# Initiation of organ maturation and fruit ripening in grapevine is controlled by the CARPO-NAC transcription factor

**DOI:** 10.1101/2021.11.13.468481

**Authors:** Erica D’Incà, Chiara Foresti, Luis Orduña, Alessandra Amato, Elodie Vandelle, Antonio Santiago, Alessandro Botton, Stefano Cazzaniga, Edoardo Bertini, Mario Pezzotti, James Giovannoni, Julia Vrebalov, José Tomás Matus, Giovanni Battista Tornielli, Sara Zenoni

## Abstract

Grapevine is a woody temperate perennial plant and one of the most important fruit crops with global relevance in both the fresh fruit and winemaking industries. Unfortunately, global warming is affecting viticulture by altering developmental transitions and fruit maturation processes. In this context, uncovering the molecular mechanisms controlling the onset and progression of ripening could prove essential to maintain high-quality grapes and wines. Through a deep inspection of previously published transcriptomic data we identified the NAC family member *VviCARPO* (Controlled Adjustment of Ripening and maturation of Plant Organs) as a key regulator of grapevine maturation whose induction precedes the expression of well-known ripening associated genes. We explored VviCARPO binding landscapes through DAP-seq and overlapped its bound genes with transcriptomics datasets from stable and transient VviCARPO overexpressing grapevine plants to define a set of high-confidence targets. Among these, we identified key molecular ripening markers. Physiological, metabolic and promoter activation analyses showed that VviCARPO induces chlorophyll degradation and anthocyanin accumulation through the up-regulation of *VviSGR1* and *VviMYBA1*, respectively, with the latter being up-regulated through a VviCARPO-VviNAC03 regulatory complex. Despite showing a closer phylogenetic relationship to senescent-related AtNAP homologues, *VviCARPO* complemented the *nor* mutant phenotype in tomato, suggesting it may have acquired a dual role as an orchestrator of both ripening- and senescence-related processes. Our data supports CARPO as a master regulator of the grapevine vegetative-to-mature phase organ transition and therefore an essential target for insuring fruit quality and environmental resilience.

**SIGNIFICANT STATEMENT:** CARPO is a grape NAC transcription factor central to fruit ripening and tissue senescence. This regulator influences multiple biological pathways common to both processes including cell wall metabolism, chlorophyll degradation, pigment production and hormone synthesis/signaling through regulation of their key genes. As various external stresses and changing climatic conditions influence vegetative growth and berry ripening, *CARPO* could prove a useful genetic and breeding target towards maintaining necessary crop performance and fruit-quality characteristics.

## INTRODUCTION

Plant lifespans are marked by ordered entry and exit to different developmental phases. Complex signaling networks, involving time-dependent interactions among hormones, transcription factors, microRNAs and environmental stimuli, govern phase transitions such as the juvenile-to-adult vegetative phases (1, 2), ensuring dynamic and fine-tuning adjustments of the plant developmental program also in the context of biotic and abiotic external factors.

At the end of aging, plant tissues undergo senescence, a finely regulated deterioration process, involving ordered physiological, biochemical and metabolic changes, crucial for recycling of materials and energy (3). Ripening is an active pre-senescence biological process during which fruit undergo dramatic changes to became attractive for animals that will promote the dispersion of seeds though ultimately terminating in tissue senescence. Senescence and ripening are both accomplished by chlorophyll degradation, secondary metabolite accumulation and cell wall breakdown and dismantling and involve at least some common signaling and regulatory factors including ethylene and transcription factors (4).

At the molecular level, it has been shown that members of the NAC (NAM, ATAF, and CUC) transcription factor family, play key roles in the regulation of leaf senescence and fruit ripening by interacting with other transcription factors, hormones and environmental signals (5, 6). Tomato NON-RIPENING (NOR) was the first NAC TF described as a master regulator of fruit ripening (7-9), and more recently shown to play a key role in leaf senescence (10).

NAC genes act as both transcriptional activators and repressors and this duality may be achieved by recruiting or interacting with different transcriptional partners (11). In peach, two NAC TFs, BL and PpNAC1, have been shown to work in a heterodimeric complex for the activation of the R2R3 MYB gene *PpMYB10*.*1*, a regulator of anthocyanin levels in fruit flesh during maturation (12). NAC TFs are also involved in ripening of non-climacteric fruits, as the NAC gene FaRIF which controls critical ripening-related processes in strawberry (13) and FcrNAC22 which mediates red light-induced fruit coloration by enhancing the expression of carotenoid biosynthetic genes in citrus (14).

Grapevine (*Vitis vinifera* L.) is one of the oldest and most widely cultivated perennial non-climacteric fruit crops in the world. Grapevine fruit development is marked by dramatic physiological, transcriptional, metabolic, and textural changes that coincide with the onset of ripening (15). The quality of grapes is extremely sensitive to environmental factors and global warming is negatively affecting the maturation process, by accelerating the onset of ripening (16, 17). Clearer understanding of the molecular mechanisms controlling plant organ development and developmental phase transitions is essential in developing strategies for optimizing the grape ripening process and related fruit quality traits in the context of climate change and additional abiotic stresses.

A clear transcriptional distinction between vegetative/green and mature/woody tissues has been shown (18) and co-expression network analyses have led to the identification of a set of candidate genes, called *switch* genes, as putative key regulators of the organ phase transition to mature growth in grapevine (19, 20). Two NAC family members, *VviNAC33* and *VviNAC60* have been described as *switches*. We recently demonstrated that VviNAC33 induces leaf de-greening through activation of the *STAY-GREEN PROTEIN 1* (*SGR1)* and growth cessation via inhibition of *AUXIN EFFLUX FALICITATOR PIN1* and *RopGEF1* expression, and proposed this NAC as a key controller of senescence-associated processes both in leaf and berry (21).

Here, we present a functional characterization of *VviNAC60*, renamed as *VviCARPO*, for the Greek goddess of Autumn and fruit harvest. We provide evidences for its role as a necessary component of fruit ripening. Through combined DNA-binding, transcriptome activity and functional analyses, *VviCARPO* high confidence promoter targets were identified, revealing its regulative role in secondary metabolism, cell death and organ de-greening. Stable transgenic lines overexpressing *VviCARPO* and its transgenic chimeric repressors showed clear alteration of senescence processes. In addition, the overexpression of the *VviCARPO* in the tomato *nor* mutant rescued the *nor* mutation at the same level as the tomato NAC-NOR. Together, these results define VviCARPO as a necessary grape ripening and senescence regulator and a target for modification/breeding toward climate resilience and fruit quality.

## RESULTS

### NAC60-CARPO represents a regulator of vegetative-to-mature phase transition in grapevine organs

*NAC60* was previously found among putative master regulators of the vegetative-to-mature phase transition in grapevine organs (19). This gene shows an increase in expression from vegetative/growth to the senescence/ripening phase in all organs, especially in all berry tissues and leaves examined (**Fig. 1A**; ***SI Appendix*, Fig. S1**).

**Fig.1.**
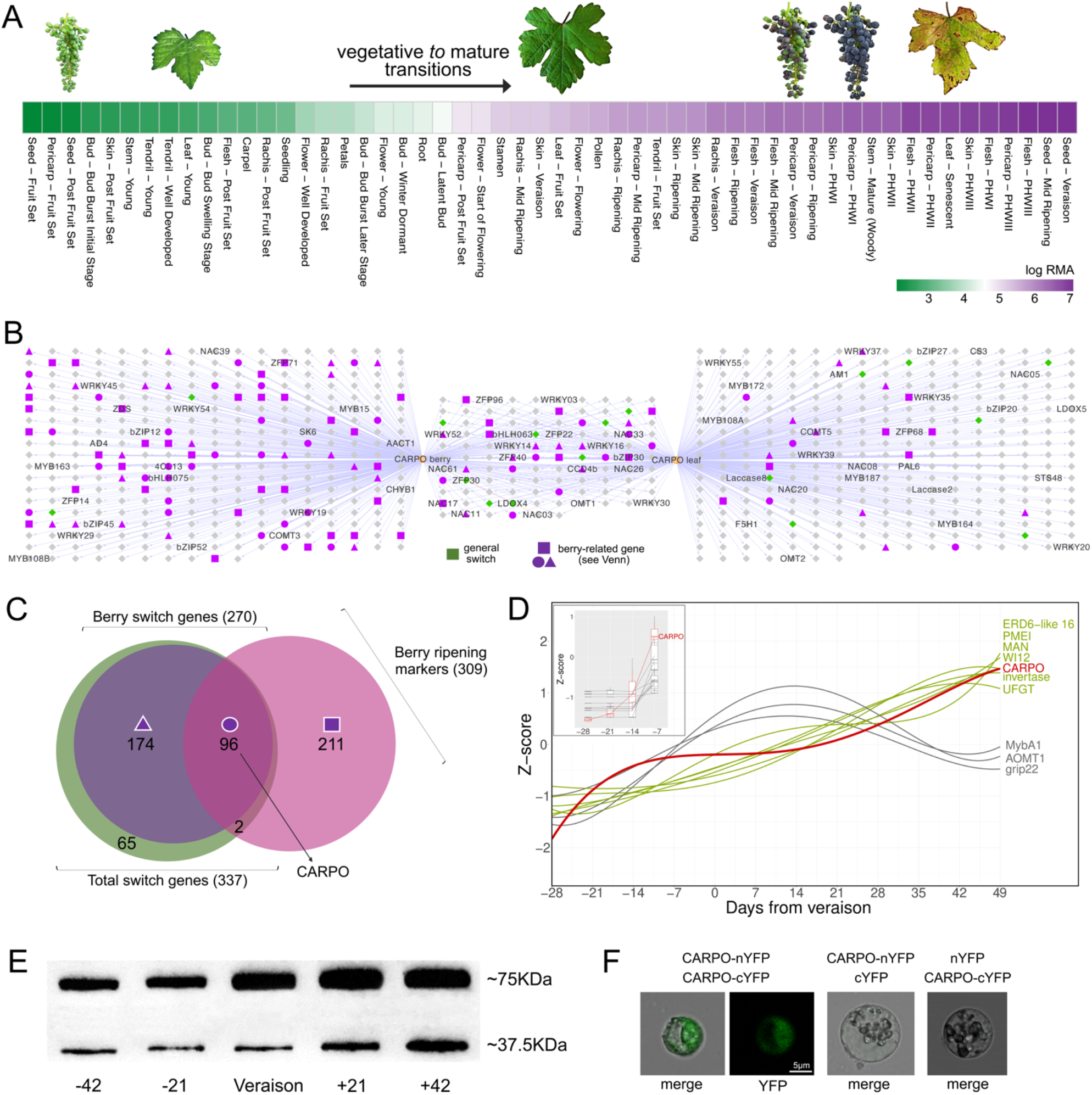
CARPO expression throughout organs development and self-interaction in grapevine. (*A*) Gene expression behavior of NAC60 (named hereafter *CARPO*) in grapevine organs throughout development. The data were sourced from the atlas transcriptomic dataset of cv. ‘Corvina’ (18). (*B*) Aggregate co-expression networks of the TOP420 most highly co-expressed genes with *CARPO* taken from berry and leaf condition-dependent analyses (22). Gene names appearing in the network were cross-referenced from the Grape Gene Reference Catalogue (Navarro-Payá et al., submitted). Node colors and shapes represent the genes classified in (*C*) resulting from the overlap of general and berry switch genes (identified in (19, 20)) and berry transition marker genes (15). (*D*) Expression profiles of *CARPO* and literature ripening-associated genes across a high-resolution series of berry developmental stages studied through RNA-seq (15). (*E*) Western blot analysis of different stages of berry development. Total protein extracts were blotted using anti-CARPO polyclonal antibody. Stages correspond to 42- and 21 days before veraison, veraison, 21- and 42 days post veraison, selected for correlation with CARPO and biomarker expression profiles. The CARPO protein has a molecular weight of ∼ 37.5KDa, which is represented by the lower bands; the upper bands (∼75 KDa) reflect the CARPO homodimer prevalence in all the samples. (*F*) Bimolecular Fluorescence Complementation (BiFC) analysis in grapevine protoplasts showing CARPO/CARPO protein interaction. Corresponding controls are also shown. Images show a representative case of YFP signal being detected in the cell nucleus, by using confocal laser scanning.

While inspecting *NAC60* gene co-expression networks (GCNs) constructed from publicly available berry or leaf RNA-seq datasets (more than 1,400 SRA runs; (22)), several switch genes (19, 20) and berry ripening transition markers (15) were found (**Fig.1B-C**), including *WRKY19* and *bHLH75* identified as markers of the first transition during berry ripening in (15)(***SI Appendix*, Dataset S1**). A high proportion of transcription factors from different families are highly co-expressed with *NAC60* with several berry related ‘markers’ found in the berry-GCN. Interestingly, many *NAC* family members were found in both GCNs; particularly interesting were *NAC11*, a berry specific switch (20), *NAC33*, recently characterized as a master regulator of senescence (21), *NAC26*, likely involved in regulating fruit and seed development (23), and *NAC61*, described as a postharvest withering-related gene (24).

Analysis of *NAC60* expression during berry development revealed that its induction starts before the expression of several well-known ripening-associated genes (**Fig. 1D**), including a berry sugar invertase, the berry anthocyanin regulator *MYBA1* and its targets *UFGT* and *AOMT1*, and the cell wall metabolism-related *GRIP22*. These data suggest that *NAC60* could be one of the main initiators of ripening. NAC60 protein accumulation throughput berry development was assessed using a specific polyclonal antibody showing its presence in all five stages surveyed, increasing from pre-ripening to fully ripe stages. Though the NAC60 monomer was detected, the NAC60 homodimer was the prevalent form (**Fig. 1E**). Moreover, a deeper inspection at the earlier stages of berry development indicates the NAC60 protein begins to accumulate around 42 days before the onset of ripening (i.e. *veraison;* ***SI Appendix*, Fig. S2**), in full correlation with the gene expression profile previously observed in the grapevine Atlas (**Fig. 1A**).

The capacity of NAC60 to interact with itself has been demonstrated by performing bimolecular fluorescence complementation (BiFC) analysis which also highlighted NAC60 homodimer localization into the nucleus (**Fig. 1F)**. In fact, NAC60 protein contains a monopartite N-terminal nuclear localization signal (PRDRKYP) **(*SI Appendix*, Fig. S3**). These data indicate that NAC60 regulatory activity is mainly achieved through the conformation of a NAC60-NAC60 complex potentially interacting with open chromatin.

We amplified the full-length coding region of *NAC60* from cultivar (cv.) ‘Syrah’ berry cDNA. The deduced amino acid sequence (335 aa) is 99% identical to the predicted ortholog in PN40024 based on its latest reference genome assembly (12X.2 assembly and VCost.v3 annotation; (25)), with only two amino acid substitutions in the highly divergent C-terminus at positions 223 and 256 (***SI Appendix*, Fig. S3**).

We constructed a full NAC family phylogenetic tree using 74 grapevine and 93 tomato sequences, adding different NAC proteins that have been functionally characterized in Arabidopsis and other plant species. We show that VviNAC60 belongs to the NAP-clade, including OsNAP (26), SlNAC1 AtNAP (27), CmNAC60 (28) and GhNAP (29), all involved in the regulation of senescence. The *VviNAC60* closest homolog is *AtANAC047*, induced upon senescence in leaf petioles (30). Moreover, in the same clade, we found *SlNAP2*, that regulates the expression of NOR (31) and *VviNAC26*. Interestingly, the NAP-clade is close to the NOR-clade, including *SlNOR-like 1* (32), *PpNAC1* that regulates phenylalanine biosynthesis in pine (33), the recently characterized *FvRIF*, involved in fruit ripening control in strawberry (13), and two grapevine NACs, *VviNAC03* and *VviNAC18* (***SI Appendix*, Fig. S4**). Based on the functional role of VviNAC60 inspected in this work we named this transcription factor as CARPO (<u>C</u>ontrolled <u>A</u>djustment of <u>R</u>ipening and maturation of <u>P</u>lant <u>O</u>rgans), aptly also the name of the Greek goddess of Autumn and harvest.

### CARPO induces the senescence phase

*CARPO* was overexpressed in grapevine cv. ‘Syrah’ in three independent transgenic lines (OX#1, OX#2 and OX#3) (***SI Appendix*, Fig. S5A and B**). The overexpression of *CARPO* led to slightly stunted growth compared to the control, due to a significant reduction in internode length **(Fig. 2A)**, reduced leaf area (c. 40%) (***SI Appendix*, Fig. S5C**) and accumulation of anthocyanin in young leaves **(Fig. 2A; *SI Appendix*, Fig. S5D**). We selected the line #1 (hereafter OX.CARPO) displaying the highest expression for further analysis. We also generated a dominant repressor version of CARPO by fusing the EAR-repression motif (SRDX) to the carboxyl terminus of the protein. Three independent transgenic lines of cv. ‘Syrah’ transformed with the *ProCARPO:CARPO-SRDX* construct were obtained (EAR#1, EAR#2 and EAR#3) (***SI Appendix*, Fig. S6A and B**). The plants in which *CARPO-SRDX* is under the control of the endogenous promoter displayed normal growth with increases in internode length (**Fig. 2A**) and leaf area (c. 35%) (***SI Appendix*, Fig. S6C**), but with similar leaf anthocyanin content as compared to the control (**Fig. 2A**). We selected the line #3 (hereafter CARPO.EAR) for further analysis.

**Fig.2.**
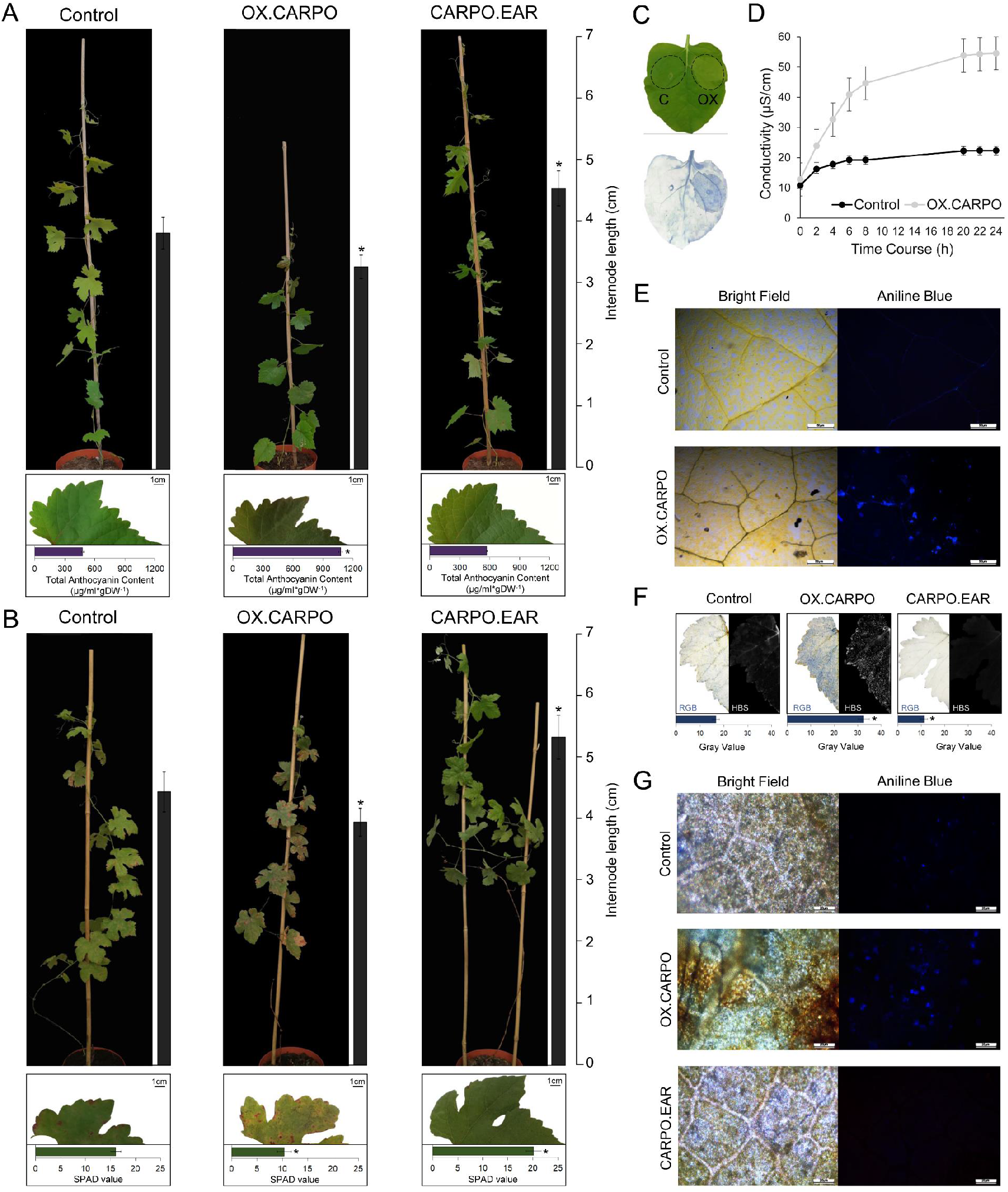
Phenotypic changes in transgenic grapevine plants with altered expression of CARPO. (*A*) Whole two months plant and leaf phenotype caused by the ectopic expression of CARPO in selected OX and EAR lines compared to vector control. Internode lengths on plants and anthocyanins contents on leaves are indicated by the bars next to each picture. The data are expressed as mean ± SD (n = 4). Asterisks indicate significant differences (*, P < 0.01; *t-*test) in the OX.CARPO lines compared to the control. SD, standard deviation. (*B*) Whole ten months plant and leaf phenotype caused by the ectopic expression of CARPO in selected OX and EAR lines compared to vector control. Internode lengths on plants and chlorophyll contents on leaves are indicated by the bars next to each picture. Asterisks indicate significant differences (*, P < 0.01; *t-*test) in the OX.CARPO and CARPO.EAR lines compared to the control. (*C*) Phenotype showed by *N. benthamiana* leaf after 72 hours from the agroinfiltration with OX.CARPO and the control (left) and cell death visualized by trypan blue staining (right). (*D*) Ion leakage assay on *N. benthamiana* leaves after 24 hours from the agroinfiltration with OX.CARPO and the control. (*E*) Callose deposition in *N. benthamiana* leaves after 72 hours from the agroinfiltration with OX.CARPO and the control visualized by aniline blue staining. (*F*) Cell death on OX and EAR CARPO grapevine leaves compared to the control visualized by trypan blue staining. The data are expressed as mean ± SD (n = 3). Asterisks indicate significant differences (*, P < 0.01; *t-*test) in the OX.CARPO and CARPO.EAR lines compared to the control. SD, standard deviation. (*G*) Callose deposition was visualized by aniline blue staining in OX and EAR grapevine leaves and the control in a bright field (on the left) and under an epifluorescence microscope (on the right). Magnification 10x.

At present, neither the CARPO transgenic plants nor the controls flower under our growing conditions, hindering our ability to determine the effects of OX.CARPO or CARPO.EAR in transgenic berries.

Ten months old OX.CARPO and CARPO.EAR transgenic plants showed opposite phenotypes in terms of symptoms of senescence compared to the control: the overexpression of *CARPO* led to premature leaf senescence, while the dominant repressor showed delayed senescence (**Fig. 2B**). Chlorophyll content was lower in OX.CARPO senescing leaves, while higher in CARPO.EAR, compared to the control (**Fig. 2B**). We further investigated the influence of CARPO on cell death in *Nicotiana benthamiana* leaves agroinfiltrated with *CARPO* under the control of a 35S promoter. At 72 hours post-infection, in contrast to the control, leaves expressing *CARPO* displayed browning necrotic regions (**Fig. 2C; *SI Appendix*, Fig. S7A**). In line with macroscopic observations, trypan blue staining (**Fig. 2C, *SI Appendix*, Fig. S7B**) and ion leakage measurement (**Fig. 2D**) clearly confirmed a significant increase of cell death when *CARPO* is ectopically expressed. Moreover, besides cell death induction, aniline blue staining revealed callose deposition in *CARPO*-agroinfected leaves (**Fig. 2E, *SI Appendix*, Fig. S7C**). Similarly, in grapevine, trypan blue and aniline blue staining highlighted cell death and callose deposition in areas showing clearer senescence symptoms in OX.CARPO and milder responses in CARPO.EAR leaves as compared to wild-type (**Fig. 2F**), consistent with the high transgene expression and the chimeric repressor activity, respectively.

### Identification of CARPO high confident targets

We carried out DNA Affinity Purification followed by Sequencing (DAP-Seq) to inspect the binding landscape of VviCARPO. Using four biological replicates and genomic DNA from leaves and berries we identified 27,715 binding events (i.e., enriched peaks compared to input), assigned to 11,310 genes, for which their position related to the transcription start site (TSS) of each gene showed a preferential distribution in proximal promoter regions (27.65% of peaks being assigned between – 3Kb and the TSS; **Fig. 3A; *SI Appendix*, Dataset S2**). Retrieving the TOP600 highest-scored peaks allowed us to identify one major binding motif (CACGTAAC) in their center. This strongly significant consensus motif was compared to Arabidopsis NAC phylogenetic binding footprints (34) showing its closest resemblance to AtANAC047’s motif, VviCARPO’s closest homolog (**Fig. 3B**; ***SI Appendix*, Fig. S4**).

**Fig.3.**
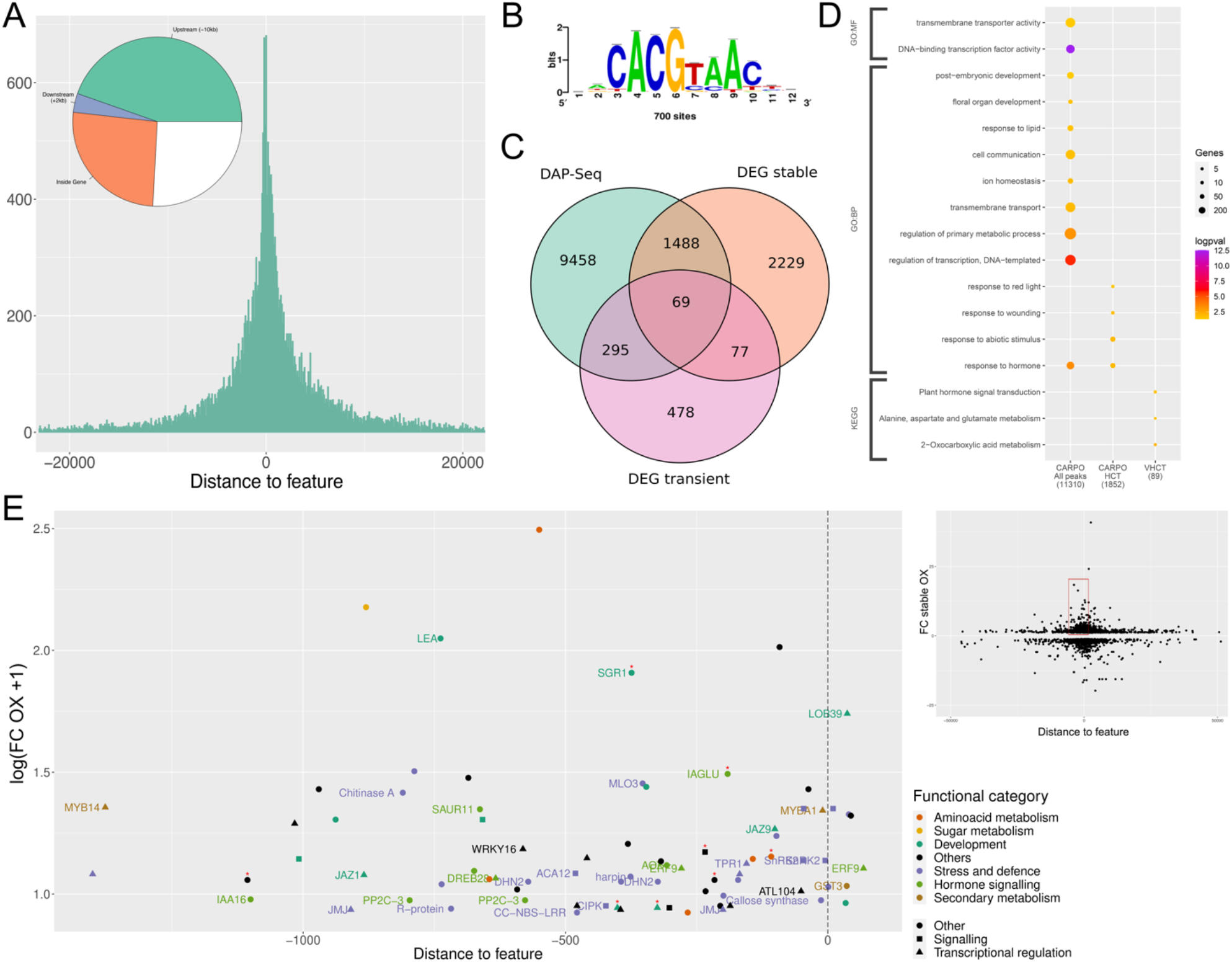
Identification of CARPO targets by DAP-Seq and transcriptomic analysis. (*A*) Distribution of CARPO DNA binding events (27,715 peaks, assigned in at least one of the three analyses: 500ng leaf, 1000ng leaf and 500ng berry) with respect to their position from the transcription start sites (TSS) of their assigned genes (n: 11,310). (*B*) *De novo* forward binding motif obtained from the inspection of the top 600-scoring peaks of CARPO 500ng leaf library using Regulatory Sequence Analysis Tools (RSAT). (*C*) Prediction of CARPO targets based on the overlap of DAP-Seq assigned genes (sum of the three analyses) and differentially expressed genes (DEGs) detected in stable and transient CARPO-overexpressing plants. (*D*) Functional enrichment analysis of CARPO bound-genes (assigned from all peaks) and selected targets (HCTs, High Confidence Targets; VHCTs: Very High Confidence Targets). VHCTs were selected by focusing on peaks in the -1.5Kb and +100bp region and on genes with FC ≥ 1.3 in either the stable or transient overexpressing lines. As additional criteria for VHCTs, we only considered peaks present at the same position from the TSS in at least two of the three analyses. Gene Ontology terms and KEGG pathways shown were filtered based on significance (Benjamini-Hochberg adjusted *p-*value < 0.05) and biological redundancy (complete list of terms can be found in **Tab. S2**). The size of each dot represents the number of genes in the input query that are annotated to the corresponding term, and the color represents the significance. Total number of genes in each list is shown in parenthesis. GO:BP, Gene Ontology: Biological Process; GO:MF, Gene Ontology: Molecular Function; KEGG, Kyoto Encyclopedia of Genes and Genomes. (*E*) Relationship between peak distances assigned to each gene and differential expression in stable and/or transient CARPO overexpression (OX) lines in selected VHCTs. Fold Change values correspond to the expression in OX lines versus the control lines, with a positive threshold of FC ≥ 1.3. Node color depicts biological processes for each gene, whereas the shape is depicted by its molecular function. Red asterisks depict those genes up-regulated in the transient overexpressing lines.

To identify CARPO’s high confident targets (HCTs), we combined our DAP-Seq data with differentially expressed genes (DEGs) induced by CARPO stable and transient overexpression. On the one hand, the transcriptomic analysis performed on young leaves of the OX.CARPO transgenic line revealed 3,863 DEGs compared with the control (P < 0.5; *t-*test; ***SI Appendix*, Dataset S3**), while the transient *CARPO* overexpression in cv. ‘Sultana’ (***SI Appendix*, Fig. S8**) identified 919 DEGs (P < 0.5; *t-*test; ***SI Appendix*, Dataset S4**). The overlap of DAP-seq data with at least one of the DEGs lists show 1,852 HCT genes (**Fig. 3C**; ***SI Appendix*, Dataset S5**), those which are mainly enriched in ‘abiotic stimuli’, ‘response to hormone’, ‘wounding’ and ‘red-light’ gene ontology categories (**Fig. 3D**). By using a more stringent criteria (e.g., excluding binding events located before –1.5kb and beyond 100bp after the TSS; see material and methods), we narrowed down our target gene set to 89 very high confident targets (VHCTs; ***SI Appendix*, Tab. S1**) genes enriched in ‘plant hormone signal transduction’, ‘amino acid metabolism’ and ‘2-oxocarboxylic acid metabolism’ descriptors (**Fig. 3D**; ***SI Appendix*, Tab. S2**; some of them being validated by qPCR, ***SI Appendix*, Fig. S9**). The expression of VHCT genes were profiled by exploring the cv. ‘Corvina’ atlas dataset revealing that most of them follows an activation pattern throughout organ development (***SI Appendix*, Fig. S10**) In particular, several genes were induced in ripening berry at higher expression levels compared to the other organs.

Among the VHCTs list we found many genes involved in plant development, such as *LATERAL ORGAN BOUNDARIES 39* (*LOB39);* in chlorophyll and photosystem degradation, such as *STAY GREEN 1* (*SGR1*; (35, 36)); in hormone signaling, such as *IAA16, IAGLU* and *SAUR11;* in secondary metabolism, such as the stilbene-related *MYB14* (22, 37) and the anthocyanin-related *MYBA1* (38), and in response to biotic and abiotic stresses, such as resistance protein-encoding genes, *HIN1* (harpin inducing protein 1-like 9), *ACA12, JAZ1, JAZ9* and *ATL104* as well as *CHITA* and a callose synthase-encoding gene corroborating the phenotype observed in *CARPO*-overexpressing plants (**Fig. 3E**).

### CARPO directly controls the expression of genes related to senescence and berry ripening

To unravel the regulatory mechanisms of VviCARPO we compared its targets with DAP-seq data generated for two other grapevine NAC family members, namely VviNAC03 and VviNAC33 (21). VviNAC03 was selected for its highly similar expression profile with VviCARPO during flowering and berry ripening (***SI Appendix*, Fig. S1**) and as the closest homolog of the tomato *SlNOR* gene (***SI Appendix*, Fig. S4**). DAP-Seq performed on VviNAC03 led to 4,277 peaks (***SI Appendix*, Dataset S6**) with the consensus sequence CACGCAAC as the most frequent binding motif (***SI Appendix*, Fig. S11A**), a footprint correlated with AtNAM and AtNAP. For VviNAC33, with a consensus motif ACA(A/C)GCAAC matching that of AtANAC017, AtNAM and AtCUC2, a total of 7,641 peaks were obtained (***SI Appendix*, Fig. S11B and Dataset S7**). We focused on the filtered DAP-Seq bound genes (i.e. with peak distances between -5Kb to +100 bp), corresponding to 4,826, 1,614 and 2,628 putative targets identified for VviCARPO, VviNAC03 and VviNAC33, respectively. All these groups of genes were mainly enriched in ‘DNA-binding transcription factor activity’, suggesting a high regulatory hierarchy for these NAC TFs. VviCARPO and VviNAC03 putative target genes were enriched in ‘plant hormone signal transduction’, ‘amino acid metabolism’ and ‘cell differentiation’, while VviCARPO specific putative target genes were enriched in ‘transmembrane transport’ and ‘response to auxin’ (**Fig. 4A**). VviCARPO showed the highest number of specific targets (2,941) and a similar number of shared genes with both VviNAC03 and VviNAC33. In contrast, fewer genes were shared between VviNAC33 and VviNAC03 (**Fig. 4A**).

**Fig.4.**
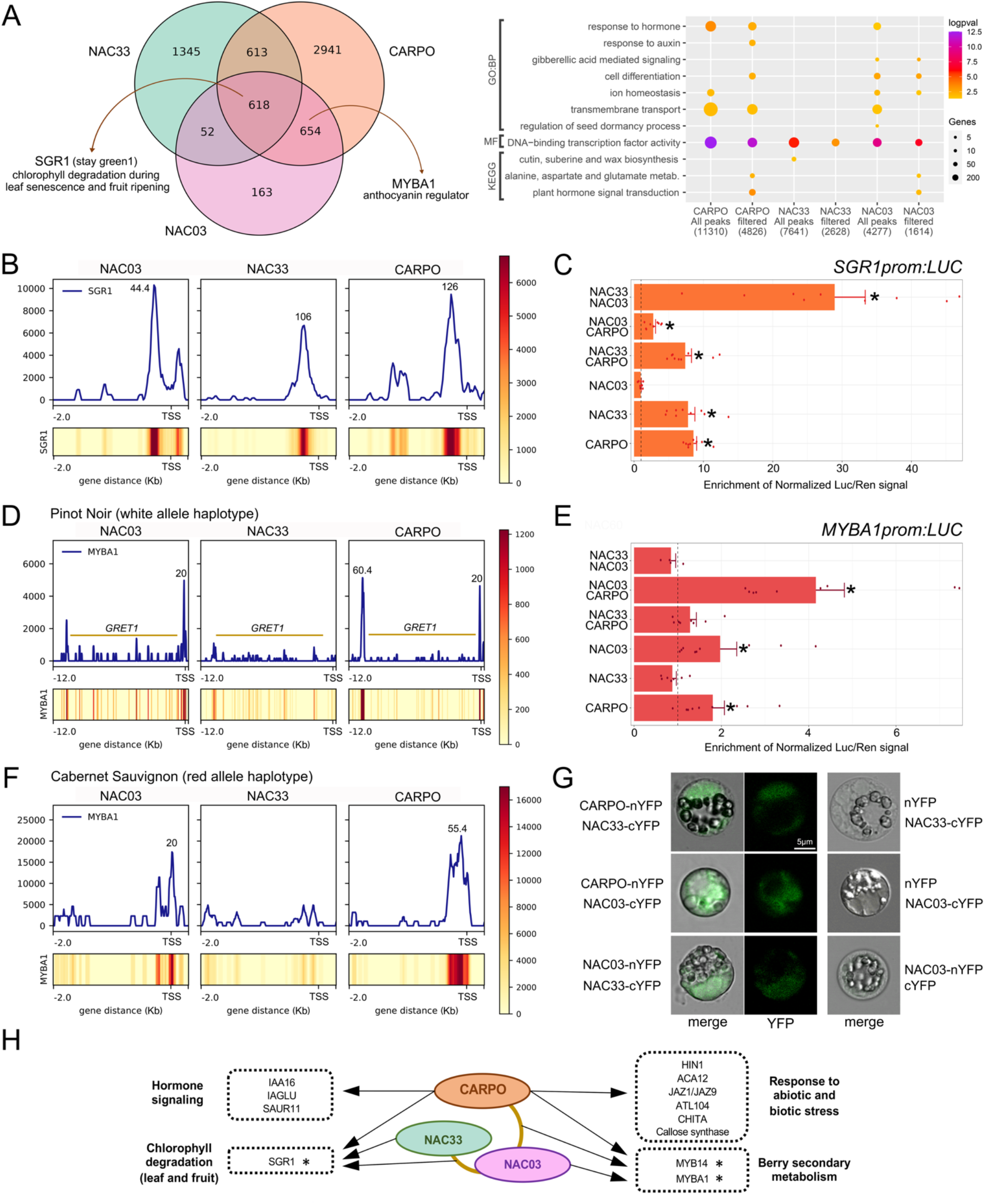
Similarities among grape CARPO and other NAC-TFs cistromes. (*A*) Overlap of NAC03, NAC33 and CARPO filtered DAP-Seq bound genes (positional filtering, with peaks ranging between -5000 bp from Transcriptional Start Site up to 100 bp from the end of the gene) (left panel); enrichment analysis of significant (P < 0.05; *t-*test) Gene Ontology terms and KEGG pathways for NAC-bound genes (right panel). (*B*) NAC DNA binding landscapes in the proximal promoter region of *SGR1* gene. DAP-Seq binding signals, shown as density plots and heatmaps and delimited between -2kb and +2kb from the TSS. Binding events identified by GEM are pointed out with their corresponding signal score. (*C*) *SGR1* promoter activation tested by dual-luciferase reporter assay in infiltrated *N. benthamian*a leaves. The single and combined activity of NAC03, NAC33 and CARPO were tested. Firefly LUCIFERASE (LUC) activity values are reported relative to the RENILLA (REN) value and normalized against the control (empty-effector vector). Each value represents the mean of three biological replicates, each with three technical replicates (± SD). Asterisks indicate significant differences in promoter activation compared with the control (*, P < 0.05; *t-*test). (*D*) NAC DNA binding events in promoter regions of *MYBA1* gene. Upper panel: Sequencing reads were mapped in the cv. ‘PN40024’ white allele haplotype that presents the insertion of the GRET1 retrotransposon in *MYBA1* proximal upstream region (histograms and heatmaps plotted between - 12kb and the TSS of the *MYBA1* gene). Bottom panel: mapping conducted in the red allele haplotype of cv. ‘Cabernet Sauvignon’ (no GRET1 insertion). (*E*) *MYBA1* promoter activation tested by dual-luciferase reporter assay in infiltrated *N. benthamiana* leaves. The single and combined activity of all NACs were tested. (*F*) BiFC analysis showing CARPO/NAC33, CARPO/NAC03 and NAC03/NAC33 protein interactions. Images are confocal laser scanning micrographs of PEG-transformed grapevine protoplasts. (*G*) Proposed regulatory mechanism model for the action of NAC-TFs on selected targets. Asterisks (*) indicate genes validated by dual-luciferase assays and the gold line indicated the interaction validated by BiFC analysis.

The comparison among the three NACs revealed 618 common target genes, supporting the hypothesis that these three NACs could participate in the regulation of common pathways. Interestingly, among them, we found *SGR1*. The binding landscape of NAC03, NAC33 and CARPO on *SGR1* promoter confirmed the binding signal of the three NACs (**Fig. 4B**). The dual-luciferase assay showed that CARPO and NAC33 significantly activated the *SGR1* promoter at the same level, while no *SGR1* promoter activity was induced by NAC03 (**Fig. 4C**). The analysis of the three combinations, CARPO/NAC33, CARPO/NAC03, NAC33/NAC03, revealed that only the latter showed a strong positive synergic effect on *SGR1* promoter activity (**Fig. 4C**).

Among bound genes in common between CARPO and NAC03 we found the anthocyanin pathway regulator *MYBA1*, one of the two main genes of the berry colour locus. As the reference PN40024 genome assembly only considers the white allele haplotype (with the presence of the *MYBA1*-inactivating GRET1 retrotransposon), we also mapped the reads from all three NAC libraries in the red allele haplotype of cv. ‘Cabernet Sauvignon’ to evaluate the effect of *GRET1* position in the binding of these TFs in *MYBA1* promoter. The analysis highlighted a very proximal binding signal for CARPO and NAC03 in both genotypes, while a further upstream binding of CARPO before the GRET1 insertion was also found in the white allele haplotype. The presence of close binding events in both CARPO and NAC03 suggests a positive interaction between these two TFs (**Fig. 4E**). No binding signal was identified for NAC33 (**Fig. 4D**). Interestingly, the binding motif recognized by CARPO on the *MYBA1* promoter diverged from the major binding motif identified in the TOP600 peaks. This consensus motif (AATGGGAC; ***SI Appendix*, Fig. S12**) was also present in the peaks identified in the promoter regions of the other berry color locus gene *MYBA2* (38) and the three *MYBA* genes from the vegetative color locus (*MYBA5, MYBA6* and *MYBA7;* (39)). We show that CARPO and NAC03, but not NAC33, significantly activate *MYBA1* promoter at the same level, and such activation further increased in presence of both TFs.

MYB14, one of the main regulators of stilbene synthesis, was found among CARPO-bound genes at -1377 bp from TSS, respectively. The dual-luciferase assay confirmed that CARPO significantly induced MYB14 expression (***SI Appendix*, Fig. S13)**.

The BiFC highlighted the formation of heterodimer combinations among the three NACs (**Fig. 4F**), suggesting that their action is exerted by an interaction with different NAC partners. The multi-spiked binding profile of CARPO in cv. ‘Cabernet Sauvignon’ *MYBA1* promoter suggests that the two peaks separated by *GRET1* are at a very close distance in the red-allele and could form part of a large binding of VViNAC03-CARPO and CARPO-CARPO hetero and homodimers, respectively.

Based in our results, we propose a model of regulatory action in which CARPO emerges as a master regulator of ripening- and senescence-related processes in grapevine by activating specific targets either alone or with the eventual contribution of its interacting partner NAC03 (**Fig. 4G**).

### CARPO is able to complement the *nor* mutation in tomato

To investigate the ability of VviCARPO to regulate fruit ripening initiation in the absence of fruit from transgenic grapevines, the gene was overexpressed in the *nor* tomato mutant. We overexpressed also *VviNAC33* and *VviNAC03*. The expression level of each transgene was measured in T_3_ fruits at breaker (Br)+7 by qPCR (***SI Appendix*, Fig. S14**). We noted a correlation between the level of transgene expression and the level of phenotypic rescue. CARPO#1, NAC03#12 and NAC33#3 lines (hereafter 35S:VviCARPO, 35S:VviNAC03 and 35S:VviNAC33, respectively) were selected for further analyses based on their higher transgene expression compared with the other lines (***SI Appendix*, Fig. S14**). 35S:VviCARPO plants showed a stunted growth in comparison to *nor* plants, confirmed by reduced internode length, while no evident alterations were observed for 35S:VviNAC03 and 35S:VviNAC33 plants (**Fig. 5A**).

**Fig.5.**
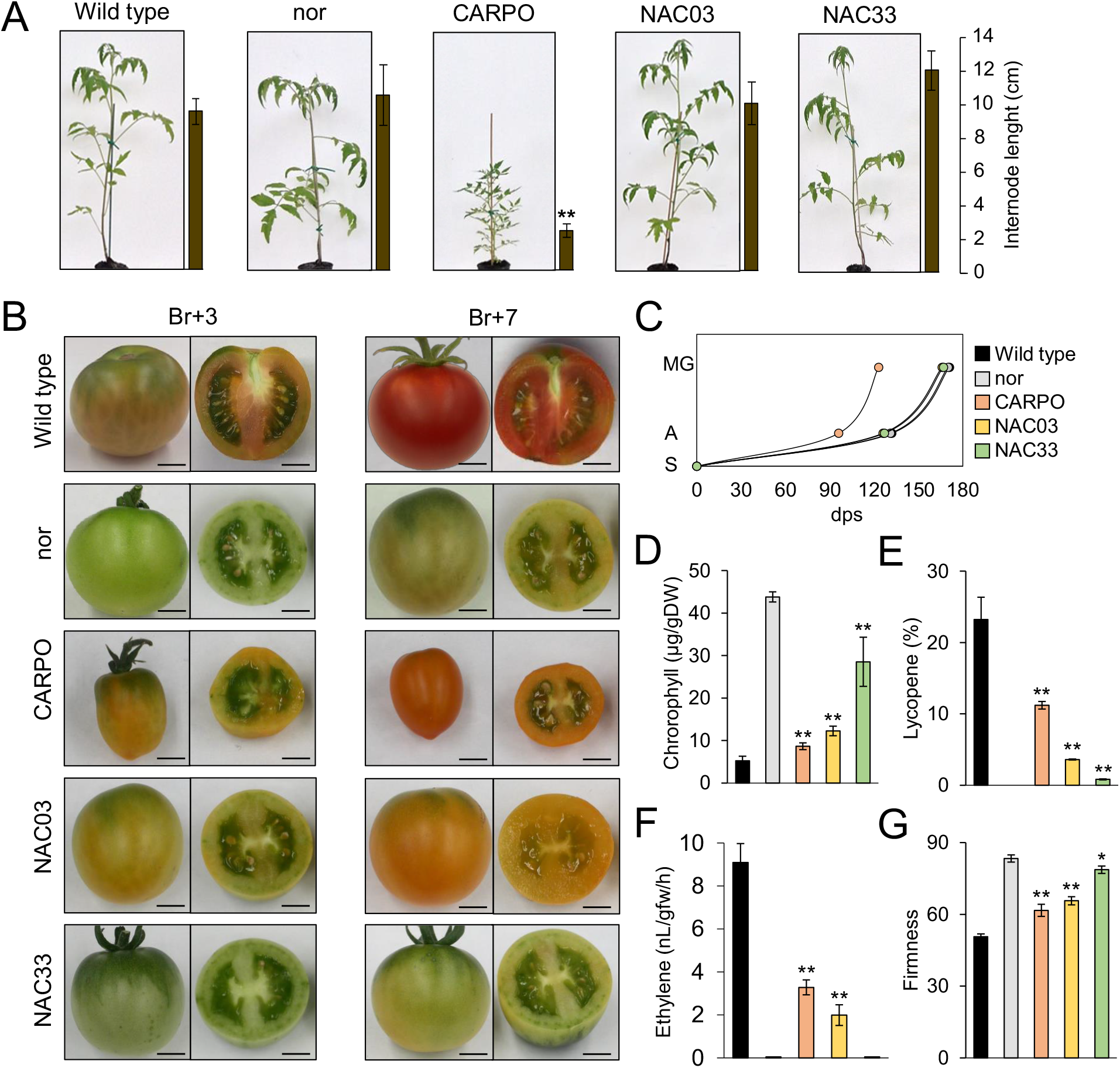
Phenotypic evaluation of VviCARPO, VviNAC03 and VviNAC33 overexpression in *nor* tomato mutant background. (*A*) Whole tomato plant phenotype corresponding to wild type (*Solanum lycopersicum* cv. ‘Ailsa Craig’), *nor* and T_3_ fruit transformed with 35S:VviCARPO, 35S:VviNAC03 and 35S:VviNAC33 in *nor* tomato mutant background. Internode lengths on plants are indicated by the bars next to each picture. (*B*) Phenotype of tomato fruits corresponding to wild type, *nor* and T_3_ fruit transformed with 35S:VviCARPO, 35S:VviNAC03 and 35S:VviNAC33 in *nor* tomato mutant background. They were collected at the breaker (Br)+3 and +7. Bar, 2cm. (*C*) Growth curve of VviCARPO, VviNAC03 and VviNAC33 transgenic plants in *nor* background, wild-type and *nor* (n = 9) from seedling to mature green stage. S, seedling; A, anthesis; MG, mature green; dps, day post seedling. (*D*) Chlorophyll content in VviCARPO, VviNAC03 and VviNAC33 transgenic fruits in *nor* background, wild-type and *nor* at Br+3. The data are expressed as mean ± SD (n = 4). Asterisks indicate significant differences (**, P < 0.01; *t-*test) compared to *nor*. (*E*) Lycopene content in VviCARPO, VviNAC03 and VviNAC33 transgenic fruits in *nor* background, wild-type and *nor* at Br+3. The data are expressed as mean ± SD (n = 4). Asterisks indicate significant differences (**, P < 0.01; *t-*test) compared to the *nor*. (*F*) Ethylene production in VviCARPO, VviNAC03 and VviNAC33 transgenic fruits in *nor* background, wild-type and *nor* at Br +5. Each value represents the mean ± standard deviation (SD) of three biological replicates. Asterisks indicate significant differences (**, P < 0.01; *t-*test) compared to *nor*. (*G*) Fruits firmness of VviCARPO, VviNAC03 and VviNAC33 transgenic fruits in *nor* background, wild-type and *nor* at Br +3. The data are expressed as mean ± SD (n = 4). Asterisks indicate significant differences (**, P < 0.01; *, P < 0.05; *t-*test) compared to *nor*.

35S:VviCARPO and 35S:VviNAC03 pericarps showed different degrees of yellowness at Br+3 with 35S:VviCARPO fruits redder than 35S:VviNAC03 at Br+7. 35S:VviNAC33 transgenic fruits were indistinguishable from *nor* in both analyzed stages (**Fig. 5B**). 35S:VviCARPO fruits were smaller than the other same-age fruits (**Fig. 5B**) and displayed earlier ripening: they flowered one month earlier than the other same-age plants and showed the interval between flowering and fruit mature green stage two-week shorter than the others lines (**Fig. 5C**).

Pigment content analysis revealed a significant decrease in chlorophyll in all transgenic fruits (**Fig. 5D**) as well as a significant increase in lycopene (**Fig. 5E**), the most abundant carotenoid in ripe tomato, in both 35S:VviCARPO and 35S:VviNAC03 in comparison to *nor*. In particular, 35S:VviNAC33 showed the highest chlorophyll content and 35S:VviCARPO the highest lycopene content, supporting the phenotypic observations.

Regarding ethylene production, we analyzed Br+5 stage since 35S:VviCARPO fruit showed the highest ethylene production at this stage (***SI Appendix*, Fig. S15**). *Nor* and 35S:VviNAC33 fruits failed to undergo an increase in ethylene production, while ethylene was significantly higher in 35S:VviCARPO and 35S:VviNAC03 fruits compared to *nor* at the same age (**Fig. 5F**). Fruit softening in 35S:VviCARPO and 35S:VviNAC03 fruits were significantly increased in comparison to same-age *nor* fruits, while no difference was measured in 35S:VviNAC33 fruits (**Fig. 5G**).

qPCR analyses performed on the 1-aminocyclopropane-1-carboxylic acid (ACC) synthase (ACS4), involved in ethylene biosynthesis, the polygalacturonase 2A (PG2A), involved in softening pathway, the phytoene synthase 1 (PSY1), involved in carotenoid pathway, and the stay-green protein 1 (SGR1), involved in chlorophyll and photosystem degradation, showed that all these genes were significantly up-regulated in 35S:VviCARPO and 35S:VviNAC03 transgenic fruits compared to *nor*, while in 35S:VviNAC33 fruits only *SGR1* was significantly up-regulated (***SI Appendix*, Fig. S16**).

## DISCUSSION

Several grapevine transcription factors have been isolated and studied in the past decade as regulators of berry metabolism and environmental and pathogen responses. Some of these characterizations have been enhanced by systems biology approaches (40, 41). However, central mediators of developmental transitions involving all these biological processes have not been identified to date. By using a wide repertoire of molecular and computational methods, we show here that the grape transcription factor NAC60, renamed as CARPO, controls different processes activated during maturation, ripening and senescence, demonstrating its role as master regulator of the vegetative-to-mature phase organ transition in grapevine.

Specially in leaves and berries, CARPO shows a clear increase from green stages to mature/senescent leaves and fully ripe fruits. Our aggregate GCN analyses in both leaves and berries revealed that *CARPO* was tightly connected with previously identified developmental transition genes (i.e. switch and ripening markers genes). In addition, *VviCARPO* was highly co-expressed with *VviNAC33*, a recently characterized master regulator of leaf senescence (21) and *VviNAC03*, the still uncharacterized closest homologue of the tomato NON-RIPENING (SlNOR) regulator (42). In this work, we found that in fact the three NACs share several putative targets and are able to form heterodimers with each other, suggesting that VviCARPO and these other two NACs cooperate in the same regulatory networks. However, several other lines of evidence support that each regulator may be particularly specialized in one or the other process, with VviNAC33 more involved in leaf senescence whereas VviCARPO, and presumably VviNAC03, in fruit ripening.

The stable *VviCARPO* overexpression in grapevine induced significant anthocyanin accumulation in young leaves by activating *VviMYBA1* expression, a positive regulator of anthocyanin synthesis in berry during ripening (38). Interestingly, *VviMYBA1* activation was further highlighted by the interaction between VviCARPO and VviNAC03. The ability of NACs to activate *MYB* genes have been demonstrated in peach, where a NAC TF, named blood (BL), activates the transcription of PpMYB10.1 by acting as a heterodimer with PpNAC1 (12) and, more recently, in apple, where MdNAC52 regulates anthocyanin biosynthesis by activating MdMYB9 and MdMYB11 (43). Here we ascertain more detail in this NAC-*to*-MYB regulation, by demonstrating that VviNAC03 binds the same *VviMYBA1* promoter region near the TSS as VviCARPO and that these two NACs are able to form a heterodimeric complex. Considering both the binding of VviCARPO and VviNAC30 to the VviMYB1 promoter and induction of VviCARPO expression with *VviMYBA1* during berry development, we propose that VviCARPO controls *VviMYBA1* activation together with VviNAC03, as a one of the first molecular events marking the onset of berry ripening.

The activation of MYB14 by CARPO further reinforces the NAC-*to*-MYB regulation scenario described by others (12). MYB14 is part of the berry late-ripening/post-ripening program (24, 44) suggesting that CARPO may cooperate with another factor(s) induced late in ripening to activate this MYB.

*CARPO* overexpression triggered senescence symptoms, such as chlorophyll degradation and cell death induction, whereas the chimeric repressors delayed senescence. We demonstrated that CARPO directly induces the expression of *SGR1*, a key regulator of leaf yellowing that interacts with chlorophyll catabolic enzymes (45), at the same level of NAC33. While CARPO and NAC33 did form a heterodimer, no additive effects were observed when analyzed in combination by luciferase assay. In contrast, *SGR1* expression was highly induced by the NAC33/NAC03 combination. Together, these results situate CARPO high in the hierarchical regulation of organ de-greening, which is a process shared during both berry ripening and leaf senescence.

CARPO-overexpressing plants accelerate the senescence program with increased cell death. Among the VHCTs identified in overlapping of DAP-seq and transcriptomics data, we found several genes involved in stress and pathogen response, including a gene encoding the *Arabidopsis Tóxicos en Levadura* (ATL) E3 ubiquitin ligase ATL104 (46). This gene belongs to a module of grapevine ATL co-expressed genes related to biotic stress (ie. CC6; (47)). This CC6 module also includes *ATL156*, whose function in grapevine resistance to *Plasmopara viticola* was recently demonstrated (48). Interestingly, *CARPO* is up-regulated in *ATL156-*overexpressing V*itis vinifera* plants. The further comparison of available datasets highlighted 13 *CARPO* targets up-regulated in *ATL156-*OE plants and 8 targets belonging to the CC6 biotic stress-related module (***SI Appendix*, Fig. S17)**. In addition, 2 CARPO targets are common to all three datasets, namely ACA12- and Harpin inducing protein 1-like 9-encoding genes, two key players in plant defense responses, in particular related to hypersensitive cell death (ref). The presence of the genes encoding RGLG1, PP2CA and SnRK2, among CARPO targets, additionally suggests a role for CARPO in ABA-mediated gene regulation. Indeed, elevated ABA inhibits the myristoylation of RGLG1 that might be translocated into the nucleus, thus facilitating its binding with PP2CA. This mechanism promotes the nuclear degradation of PP2CA and the release of SnRK2 for activating ABA responsive transcription factors. Interestingly, ABA might promote plant defense by increasing callose deposition (49) and, accordingly, a gene encoding a callose synthase was also found among the high confident targets of the transcription factor and callose deposition was demonstrated in CARPO-OX plants. Based on these observations we speculate that CARPO may also play a role in grapevine defense implementation as the cell death observed with overexpressed CARPO may be related to the hypersensitive cell death encountered in the resistance response to pathogen infection. Given the key role of ABA in triggering grape ripening (50) the presence of ABA-related genes among VHCTs further supports the involvement of CARPO in ripening control.

We observed a reduction in leaf size when CARPO is overexpressed, both in grapevine and the tomato *nor* mutant, and an increase in leaf size in response to chimeric repressor expression in grapevine. The same phenotypic effect was observed in transgenic grapevines with altered expression of NAC33, which affected the expression of genes involved in auxin metabolism (21). We found that putative targets of CARPO were mainly enriched in ‘response to auxin’ (**Fig. 4A**) and that many auxin-related genes were down-regulated in both transient and stable CARPO transgenic grapevines (***SI Appendix*, Dataset S3, S4**), strongly supporting that CARPO is involved in the control of the cessation of organ growth by affecting auxin metabolism.

Functional studies regarding fruit development and their related genes in grapevine are hindered by the difficulty to obtain berries from transgenic plant that cannot be grown in the field due to GMO legal restrictions. In fact, at present, neither the VviCARPO transgenic plants nor the controls flower under greenhouse conditions, hindering our ability to determine the effects of OX.CARPO or CARPO.EAR in transgenic berries. In this context the use of the heterologous tomato (also a berry) system represented an obvious choice for assessing VviCARPO function in fruit. We showed that VviCARPO rescues the *nor* maturation process in tomato exactly reflecting the activity of SlNOR when overexpressed in *nor* mutant (32). Interestingly, 35S:VviCARPO fruits showed a smaller dimension than control same-age fruits, as reported for 35S:NAC-NOR. Also, VviCARPO activated key genes involved in tomato fruit ripening, such as *SlACS4, SlPG2a, SlPSY1* and *SlSGR1*, though their expression level did not reach the same level as in wild-type. In *nor* fruits, a truncated NOR protein is still present and can bind to the promoters of the target genes, even though it is unable to activate them (32) and may compete with transgenic VviCARPO consistent with lower target gene expression. The complementation analysis also showed that *VviNAC03*, the closest homolog of *SlNOR*, is also involved in ripening progression control, being able to activate the previously mentioned tomato ripening genes, albeit at a lower level than VviCARPO. The overexpression of *VviNAC33* in *nor* was not able to rescue ripening, though significant chlorophyll degradation was observed, confirming its role in organ de-greening.

Overall, we showed that CARPO is a master regulator of organ phase transition by activating secondary metabolism, chlorophyll degradation, hormone metabolism, defense and cell death-related genes. Its participation to the regulation of berry ripening is inferred by the following evidence: i) it’s mRNA is strongly induced at the onset of berry ripening and correlated with expression of known ripening ‘marker’ genes; ii) a large part of its VHCTs identified by DAP-seq and transcriptomic analyses are highly expressed in ripening berry; iii) it binds and directly activates *SGR1, MYBA1, MYB14* genes associated with ripening onset and progression; iv) it complements the tomato *nor* mutation similarly to the endogenous NAC *SlNOR* gene.

While additional studies describing time-evolving regulatory networks will be necessary to better clarify the regulatory interactions of CARPO and other grapevine NACs, the functional characterization reported here defines CARPO as a key component of the molecular mechanisms governing ripening and senescence in grapevine. In the context of global climate change, uncovering and modifying the molecular components of ripening will be essential to maintaining high quality grapes and wine. This may include altering the duration of the vegetative growth phase to modify the timing of the onset of ripening. This will help to reverse the current trend towards earlier ripening and will thus avoid the harvesting of grapes under unfavorable temperature conditions, an emerging problem leading to poor-quality grape products.

## MATERIALS AND METHODS

### Plant Material, Bioinformatics, Metabolite and Molecular Analysis

Details of plant growing conditions and sample collection, molecular analyses (DNA constructs, grapevine and tomato transformation, RNA-seq, DAP-seq library construction, qPCR, protein gel-blot analysis, dual-luciferase/BiFC assays), bioinformatics procedures (gene co-expression network construction, DAP-seq and transcriptomic analyses), metabolite quantification (ethylene and pigments) and physiological measurements (fruit firmness, electrolyte leakage assay and callose staining) are provided in ***SI Appendix***.

All primers used in this work are listed in ***SI Appendix*, Table S3**.

### Data availability

Microarray data for the transient expression experiments are available at GEO under the accession GSE185447. Microarray data for the transgenic plants are available at GEO under the accession GSE185448. DAP-seq raw data are available at GEO under the accession GSE186656, including metadata of samples and conducted analysis (bioinformatic parameters) according to the FAIR principles. DAP-seq results on CARPO, NAC03 and NAC33 can be visualized in the DAPBrowse tool available at the Vitis Visualization Platform (https://www.tomsbiolab.com/vitviz).

## ACKNOWLEDGMENTS AND FUNDING SOURCES

This work was supported by Grant Ricerca di Base ‘Definition of master regulator genes of fruit ripening in grapevine’, University of Verona, awarded to SZ, by PRIN 2017 “Regulation of gene expression in grapevine: analysis of genetic and epigenetic determinants” to MP, by National Science Foundation grant IOS-1855585 and the United States Department of Agriculture - Agricultural Research Service to JJG and by grants PGC2018-099449-A-I00, RYC-2017-23645 and PRE2019-088044 to JTM and LO from the Ministerio de Ciencia, Innovación y Universidades (MCIU, Spain), Agencia Estatal de Investigación (AEI, Spain), and Fondo Europeo de Desarrollo Regional (FEDER, European Union). This article is based upon work from COST Action CA 17111 INTEGRAPE, supported by COST (European Cooperation in Science and Technology).

