## Supplemental Methods and Figures for "Initiation of organ maturation and fruit ripening in grapevine is controlled by the CARPO-NAC transcription factor"

This PDF file includes:
SI Materials and Methods
Figures S1 to S19
Tables S1 to S4
SI References

Other supplementary materials for this manuscript include the following:
Datasets S1 to S9

#### SI Materials and Methods

##### Plant material

*Nicotiana benthamiana* (*N. benthamiana*) plants, *Vitis vinifera* (*V. vinifera*) cv. Sultana plantlets, cv. 'Syrah' embryogenic calli for the genetic transformation, *VviCARPO* and *VviCARPO-EAR* transgenic plant*s* were grown as described in ref (1). For the DAP-seq analysis, cv. 'Syrah' fruiting cuttings were propagated as described in ref (2).

Embryogenic calli of *V. vinifera* cv. 'Sultana' for protoplast isolation and transfection were initiated in the 2016 season from leaf disks as described by ref. (3). Embryogenic calli were maintained and subcultured according to ref (4).

Wild-type (WT) tomato (*Solanum lycopersicum*) cv. Ailsa Craig (AC), *nor* mutant and 35S:VviCARPO, 35S:VviNAC03 and 35S:VviNAC33 plants were grown under controlled greenhouse condition with natural light. Flowers were tagged at anthesis to record the ripening stages accurately through fruit development. Fruits harvested and collected at different ripening stages: mature green (MG), breaker (Br), and 3, 5 and 7 days after breaker (Br+3, Br+5 and B+7, respectively). Pericarp tissues of the harvested fruits were collected and frozen in liquid nitrogen immediately and stored at -80 °C until use.

##### Gene co-expression network construction

Berry and leaf GCNs were produced as in (5). A total of 35 and 42 Transcriptomic RNA-Seq Sequence Read Archive (SRA) studies were explored, encompassing 807 and 670 runs from fruit/fw and leaf samples, respectively. Reads were trimmed with fastp (6), version 0.20.0 with the following parameters: "–detect_adapter_for_pe –n_base_limit 5 cut_front_window_size 1 cut_front_mean_quality 30 –cut_front cut_tail_window_size 1 cut_tail_mean_quality 30 –cut_tail -l 20". After the trimming, the runs are aligned with STAR (Dobin et al., 2012), version 2.7.3a, using default parameters. Raw counts were computed using FeatureCounts (7), version 2.0.0 with the gene models of the VCost.v3_27.gff3 available at https://integrape.eu/resources/genes-genomes/genome-accessions/. Each SRA study was individually computed to build a highest reciprocal rank (HRR) matrix (8). Each raw counts matrix was normalized to FPKMs, and genes with less than 0.5 FPKMs in every run of the SRA study were removed. The Pearson's correlation coefficients (PCC) of each gene against the remaining genes was then calculated for each SRA study (across all run matrices) and ranked in descending order. Ranked PCC values were used to compute HRRs amongst the top 420 ranked genes (420 roughly equals 1% of all VCost.v3 gene models), using the following formula: HRR(A, B) = max(rank(A,B), rank(B,A)), generating a HRR matrix for each SRA study. To construct the aggregate whole genome co-expression network, the frequency of co-expression interaction(s) across individual HRR matrices was used as edge weights, and after ranking in descending order, the top 420 frequency values for each gene were chosen to build the final aggregate networks. The top 420 most highly co-expressed genes for CARPO was used to generate its gene-centred co-expression networks (GCNs). The expression profiles of grapevine NAC genes were analyzed across 807 transcriptomic datasets (i.e., SRA runs, used for generating the GCNs) that were manually classified in three tissue categories (veraison/post-veraison berry, pre-veraison berry and inflorescence/flower). This data has been integrated in the EXHARA tool available in the Vitis Visualization Platform (VitViz; available at https://tomsbiolab.com/vitviz).

**Phylogenetic analysis**

The complete protein sequences of previously-characterized NAC family genes were used for phylogenetic reconstruction. Protein alignments were performed in MEGA X (9) applying the MUSCLE algorithm (default parameters). The best substitution protein model (with the lowest Bayesian information criterion -BIC- score) was identified using MEGAX, corresponding to 'Jones-Taylor-Thornton (JTT)+G+F'. The phylogenetic analysis was then conducted by bayesian inference, using the Markov Chain Monte Carlo algorithm (MCMC) available in MrBayes (10), using 4 runs and chains, and 1,500,000 generations obtaining an average standard deviation of split frequencies of 0.077245. The different clusters' reliability were calculated with the posterior probabilities, being the values higher than 0.70 acceptable and the values higher than 0.90 highly supported. Protein sequences are available in ***SI Appendix***, **Dataset S9.**

**Isolation and cloning**

The *VviCARPO* coding sequence (with 3′ UTR) and *VviCARPO-EAR* were amplified from cv. Syrah mid-ripening and ripening berry skin, the *VviNAC03* coding sequence (with 3′ UTR) was amplified from cv. Corvina mid-ripening (11) and the *VviNAC33* coding sequence (with 3'UTR) was amplified as described by ref. (12). All the amplifications were performed using KAPA HiFi DNA polymerase (KAPA Biosystems, Wilmington, MA, USA) and the primers listed in ***SI Appendix***, **Table S3**. The PCR products were directionally cloned into the Gateway entry vector pENTR/D-TOPO (Invitrogen, Thermo Fisher Scientific, Waltham, MA, USA).

For agroinfiltration in *N. benthamiana* and the transient expression, the *VviCARPO* sequence was transferred into the binary overexpression vector pK7GW2.0 (Laboratory of Plant Systems Biology, Ghent University, Belgium) by site-specific LR recombination. For stable overexpression, the sequence was transferred to the modified binary vector pK7GW2.0 containing an eGFP expression cassette driven by the Arabidopsis ubiquitin 10 promoter. The *VviCARPO-EAR* chimeric repressor was transferred into the modified pK7WG2 vector, harboring the 1093-bp endogenous *VviCARPO* promoter instead of *the cauliflower mosaic virus (CaMV) 35S* promoter. The *VviCARPO* putative regulative regions were amplified by Nested PCR from cv. Syrah genomic DNA by using KAPA HiFi DNA polymerase and primers listed in ***SI Appendix***, **Table S3.**

**llumina genomic DNA libraries preparation**

The Illumina genomic DNA libraries were prepared as described in ref. (12). Briefly, 7500 ng of *V. vinifera* cv. Syrah purified genomic DNA from young leaves or green berries was diluted in elution buffer (EB, 10 mM Tris HCl, pH 8.5) sheared to 200 bp fragments using a Covaris S2 sonicator. The fragments size selection was obtained using AMPure XP beads (2:1 beads to DNA ratio) and the eluted DNA was end repaired using the End-It kit (Lucigen). End-repaired DNA was purified using the Qiagen QIAquick PCR purification protocol and A tailed using Klenow 3′-5′ exo- (NEB). A-tailed DNA was purified using the Qiagen QIAquick PCR purification protocol and the purified DNA was ligated to truncated Illumina adapters overnight at 16 °C using T4 DNA ligase (NEB). The adapter-ligated libraries were purified using AMPure XP size selection beads at a 1:1 beads to DNA ratio.

**Transcription factors *in vitro* translation**

The *VviCARPO* and *VviNAC03* sequences were recombined from the pENTR/D-TOPO entry vector to the Gateway-compatible pIX-HALO destination vector (13) using LR Clonase II (Life Technologies). The HALO-VviCARPO, HALO-VviNAC03 and GST-HALO (used as negative control) fusion proteins were *in vitro* translated using 1000 ng of pIX-HALO-TF plasmid in three different TNT$^R$ SP6 coupled reticulocyte lysate system (Promega) reactions.

**DAP-seq**

The DAP-seq was performed as described in ref. (12) and a total of 3.5 million reads were obtained for each sample. In brief, the *in vitro* translated proteins were incubated with the Magne-HALOTag beads (Promega) for 1 h at room temperature in 1× PBS with 0.005% Nonidet P-40 (PBST). Bound proteins were six times washed with PBST and incubated with the adapter-ligated genomic DNA libraries for 1 h at room temperature. Beads were washed again with PBST and resuspended in EB; the DNA were then eluted heating the samples at 98 °C for 10 min. The obtained DNA were transferred to a new tube and PCR enriched as described in ref. (12) prior to sequencing on an Illumina NextSeq 500 using 75-bp single-end reads.

Four DAP-seq replicates results were respectively analyzed to study the VviCARPO and VviNAC33 cistrome: three replicates were obtained putting the *in vitro* translated transcription factors in contact with the young leaves adapter-ligated genomic DNA libraries (one using 1000 ng and two using 500 ng) and one replicate was obtained using 500 ng of green berries adapter-ligated genomic DNA libraries. Only one replicate was analyzed for the VviNAC03 cistrome analysis, which was obtained putting 500 ng of young leaves adapter-ligated genomic DNA libraries in contact with the *in vitro* translated transcription factor.

**Details on DAP-seq analysis (read mapping, filtering, peak calling and motif analysis).**

TF-associated peaks were identified as in (5). DAP-Seq reads were mapped to the 'PN40024' 12X.v2 reference genome using bowtie2 (14) version 2.0-beta7 with default parameters and post-processing to remove reads that have MAPQ scores lower than 30. Peak detection was performed using GEM peak caller (15) version 3.4 with the 12X.v2 genome assembly using the following parameters: "–q 1 –t 1 –k_min 6 –kmax 20 –k seqs 600 –k_neg_dinu_shuffle", limited to nuclear chromosomes. The biological replicates were analysed as multi-replicates with the GEM replicate mode. Peak summits called by GEM were associated with the closest gene model in the custom annotation file using the BioConductor package ChIPpeakAnno (16) with default parameters (i.e., NearestLocation). Metagene plots were produced using Deeptools suite v.3.3.2, computing and normalizing the coverage for each BAM file with a BinSize = 10 and RPKM normalisation. RPKM value for each bin is the average between the 10 positions that define each bin. BigWig files for the individual replicates for each TF were merged using bigWigMerge v.2 and bedGraphToBigWig v.4. *De novo* motif discovery was performed by retrieving 200 bp sequences, centered at GEM-identified binding events, for the 600 most enriched peaks for each TF and running the RSAT software (http://rsat.eead.csic.es/plants/peak-motifs_form.cgi), with default parameters.

**Total protein extraction and quantification**

Total protein extracts were extracted from *V. vinifera* cv. Syrah berries as described in ref. (17). The six berry samples were harvested at 49-, 42- and 21- days before veraison, at veraison, 21- and 42- days post veraison. The total protein extracts were quantified using the Bradford Reagent (SIGMA) according to the manufacturer's recommendations.

**Western blot analysis**

40 µg of total protein extracts were mixed with SB4X (40% sucrose, 8% SDS,0.25M Tris HCl pH 6.8, bromophenol blue) and heated at 100°C for 10 minutes. The samples and the ladder (PageRuler™ Prestained Protein Ladder 180KDa) were then loaded on 12% SurePAGE Bis-Tris 10-well gel (GenScript) and the gel was run at 160 volts in Tris-MOPS-SDS Running Buffer (GenScript) until a good separation of pre-stained standards was obtained. The proteins were transferred from the gel to the membrane using the Trans-Blot® Turbo™ RTA Transfer Kit (BioRad) and the membrane was blocked for 2 hours at RT in 4% milk-PBS 1X. After the saturation, the membrane was incubated

overnight at 4°C with the primary anti-VviCARPO antibody (polyclonal antibody, used 1:10000 in 4% milk-PBS 1X-0,1% Tween20) and then washed 3 times for 10 minutes in 4% milk-PBS 1X-0,1% Tween20. The incubation with the secondary anti-rabbit antibody (SIGMA; used 1:10000 in 4% milk-PBS 1X-0,1% Tween20) was performed for 2 hours at room temperature and the membrane was then washed 3 times for 10 minutes in 4% milk-PBS 1X-0,1% Tween20. At the end, the chemiluminescent detection was performed with the ECL Western Blotting substrate (Promega). The anti-VviCARPO customised polyclonal antibody was produced by Biotem company immunizing New Zealand White Rabbits with two peptides (KPPLKLRDASMRLDNWVLC and MENQLKRQRSSMDGDVPC), which were selected as the most VviCARPO specific between the VviNAC family members.

**Transient expression**

For *N. benthamiana*, pK7WG2.0 vectors containing 35S:*VviCARPO* or a non-coding sequence (negative control) were transferred to *Agrobacterium tumefaciens* (*A. tumefaciens* ) strain C58C1 by electroporation (18). Three fully-expanded leaves were syringe infiltrated and the phenotypic analysis was carried out 3 days after agroinfiltration.

The same vectors were used in 5-week-old *in vitro* plantlets of cv. Sultana (seven plants for *VviCARPO* overexpression and five for the control) as previously described (19). The material for RNA extraction and transcriptomic analysis was collected 7 days after the agroinfiltration.

**Transgenic plants**

In grapevine, the pK7WG2.0 vectors containing 35S:*VviCARPO* or *VviCARPO-EAR* or the eGFP sequence (negative control) were transferred to *A. tumefaciens* strain EHA105 by electroporation (18). The genetic transformations were performed as described in ref. (1).

In tomato, the pK7WG2.0 vectors containing 35S:*VviCARPO,* 35S:*VviNAC03* and 35S:*VviNAC33* were transferred to *A. tumefaciens* strain LBA4404 by electroporation (18).

Transformation of *nor* mutant tomato cv. AC cotyledon explants was performed at the Boyce Thompson Institute transformation facility as described by ref. (20). $T_0$ plants were grown to maturity and two independent lines for each transgene was selected by phenotype (*VviCARPO #1-#3; VviNAC03#1-#12; VviNAC33#2-#3*). They were characterized by a slight pericarp pigmentation, except for *VviCARPO#1* line that showed a reddish fruit surface, in comparison to the *nor*, that appeared completely green (***SI Appendix,*** **Fig. S18**). Seeds obtained from the two selected $T_0$ lines were planted and 76 $T_1$ plants were grown to maturity in the greenhouse (***SI Appendix,*** **Tab. S4**). The expression level of each transgene was measured in five plants per line by quantitative polymerase chain reaction (qPCR) in leaves (***SI Appendix,*** **Fig. S19**). The plants with the highest expression level for each transgene for each line were selfed to produce $T_2$ progeny and subsequently the same procedure was used to obtain $T_3$ generation. The analyses were performed on $T_3$ generation.

**Internode length estimation**

Internode length was determined by measuring with a ruler (cm) the length between every node, that is between the beginning of the first fork in the stem until the first leaf below the last inflorescence. Mean ± Standard Deviation (SD) of four biological replicates (each the average of three technical replicates) was calculated for each genotype in grapevine and in tomato plants.

**Leaf area measurement**

Leaf area was estimated as proposed by ref. (21) by measuring maximum length and width of three laminae of mature leaves (at the $6^{th}/7^{th}$ node from apex). Mean ± Standard Deviation (SD) of 12 biological replicates (each the average of three technical replicates) was calculated for each genotype.

**SPAD Measurements**

A SPAD-502 Plus Chlorophyll meter R (Konica Minolta) was used to acquire SPAD values of the fully expanded leaves of the old OX.CARPO, CARPO.EAR and WT plants. Four plants for each genotype were analyzed and values from eight leaves per each plant were sampled and averaged to a single SPAD value.

**Pigment analysis**

Chlorophyll content from grapevine leaves and tomato fruits were measured according to (22). Carotenoid content was analyzed by high- performance liquid chromatography (HPLC) upon pigment extraction in 80% acetone as described in ref. (23). Anthocyanins were extracted from grapevine leaves and analyzed according by ref. (23). Mean ± Standard Deviation (SD) of four biological replicates (each the average of three technical replicates) was calculated for each genotype.

**Ethylene measurement**

Tomato fruits of WT, *nor,* VviCARPO, VviNAC03 and VviNAC33at Br+5 were harvested, weighed and placed at room temperature for 3 h to avoid measuring ‘wound ethylene’. Subsequently, each fruit was transferred into 300 ml gas-tight jars, tabbed, sealed, and incubated at room temperature. After 4 h, 1 ml gas samples were withdrawn immediately and then analyzed by a gas chromatography equipped with a flame ionization detector. Ethylene concentrations were calculated by comparing sample peak areas with ethylene standards of known concentration and normalized for fruit weight.

Mean ± Standard Deviation (SD) of three biological replicates (each the average of three technical replicates) was calculated for each sample.

**Firmness measurement**

The fruit firmness of WT, *nor,* VviCARPO, VviNAC03 and VviNAC33 were measured at Br+3 by durometer hardness tester. In this system the larger the number, the firmer the fruit. Each fruit was measured at two or three sites. The data for each fruit were averaged as one biological replicate and each measurement was performed using three biological repetitions.

**Quantitative PCR (qPCR)**

Total RNA was isolated from grapevine apical leaves (for transient expression), fully expanded leaves (for OX.CARPO transgenic plants overexpressing *VviCARPO* and for CARPO.EAR transgenic plants expressing the chimeric repressor) and from tomato leaves and fruits pericarp using the Spectrum Plant Total RNA kit. Gene expression analysis by qPCR was performed as previously described (24, 25) using the primers listed in ***SI Appendix***, **Table S3**. Each value corresponds to the mean ± SD of three technical replicates relative to the VviUBIQUITIN1 (VIT_16s0098g01190) and the ACTIN (Solyc03g078400) internal controls in grapevine and tomato, respectively.

**Transcriptomic analysis**

The microarray analysis was performed with the RNA isolated for qPCR. For transient expression, the four most highly overexpressing plants (nos. 1, 2, 6 and 7) were selected and used as biological replicates, while for OX.CARPO transgenic lines, the OX1, 2 and 3 lines were used as biological replicates. The cDNA synthesis, labelling, hybridization and washing were performed according to the Agilent Microarray-Based Gene Expression Analysis Guide (v.6.5). Each sample was hybridized to an Agilent custom microarray four-pack 44K format (Agilent Sure Print HD 4X44K 60-mer; cat. no. G2514F-048771; (25) and scanned using an Agilent Scanner (G2565CA; Agilent Technologies, Santa Clara, CA, USA). Feature extraction and statistical analysis of the microarray data was conducted as reported by ref. (19). Differentially expressed genes (DEGs) were identified by Student's *t*-test ($\alpha = 0.05$), assuming equal variance among samples, and selected by fold change |1.3|).

**Dual Luciferase Reporter Assay**

The *VviMYBA1* (1237bp) and *VviMyb14* (1569 bp) putative regulative regions were amplified by Nested PCR from cv. Syrah genomic DNA by using KAPA HiFi DNA polymerase and primers listed in ***SI Appendix***, **Table S3.** Purified PCR product was cloned into the pPGWL7.0 reporter vector (http://www.vib.be/en/research/ services/Pages/Gateway-Services.aspx) to control the firefly luciferase gene (LUC). The *VviMYBA1pro:LUC, VviMyb14pro:LUC* and *VviSGR1pro:LUC* (previously engineered by ref. (12)) pK7WG2 reporter vectors, the 35S:*VviCARPO*, 35S:*VviNAC33*, 35S:*ViNAC03* effector vectors, and the *Renilla reniformis* (*R. reniformis*) reference vector (previously engineered by ref. (1)) were transferred to *A. tumefaciens* strain C58C1 by electroporation (18). Dual Luciferase Reporter Assay was carried out in *N. benthamiana* leaves as described (26, 27). Firefly and *R. reniformis* luminescence were detected using a Tecan Infinite ® M200 PLEX instrument.

**Grapevine protoplast transfection and BiFC (Bimolecular Fluorescence Complementation) analysis**

VviCARPO, VviNAC33, VviNAC03 coding sequences were cloned into the pGWcY Gateway vector. VviCARPO and VviNAC030 coding sequences were also cloned into the pnYGW gateway vector. As a positive control, the nuclear homodimerization of AtbZIP63 was tested (28) by using the pUC-SPYNEGW and pUC-SPYCEGW vectors. *V. vinifera* cv. Sultana protoplasts were isolated from embryogenic calli and transfected as described by ref. (3), cultured into multi-well plates in the dark at 25°C and analysed 1 day after transfection. The YFP signal was detected using a Leica TCS SP5 AOBS confocal microscope.

**Electrolyte Leakage Assay**

Electrolyte Leakage Assay was performed to quantify cell death. Leaf discs (~5 mm in diameter) were collected from the *N. benthamiana* leaves overexpressing 35S:VviCARPO plus control and immersed in 50 mL of non-ionic, double-distilled water. Three biological replicates, each with three technical replicates, were performed for each sample. After incubation at room temperature for 1 h with shaking at 160 rpm, conductivity of the solution was measured using a conductivity meter (Horiba scientific, Edison, NJ, USA) according to ref. (29).

**Trypan Blue Staining**

*N. Benthamiana* leaves agroinfiltrated with 35S:VviCARPO plus control and grapevine leaves from OX.CARPO lines plus control were sampled for a total of three biological replicates, each with three technical replicates. The staining was performed as described by ref. (30). The color intensity was calculated by image J software as described by ref. (31).

**Aniline Blue Staining**

Leaf discs from *N. Benthamiana* leaves agroinfiltrated with 35S:VviCARPO plus control and grapevine leaves from OX.CARPO lines plus control were sampled for a total of three biological replicates, each with three technical replicates. The material was bleached by acetone 90% at -20 ºC for 30min and then by EtOH 70% overnight. Clarified material was mounted on a microscope slide, washed by soaking in phosphate buffer 0.07M pH 7.0 for 10-30 min and then stained by soaking in 0.05% aniline blue in phosphate buffer 0.07M pH 7.0 for 40 min and observed under a Leica DM2500 fluorescence microscope.

#### Supplemental Figures

##### Fig. S1

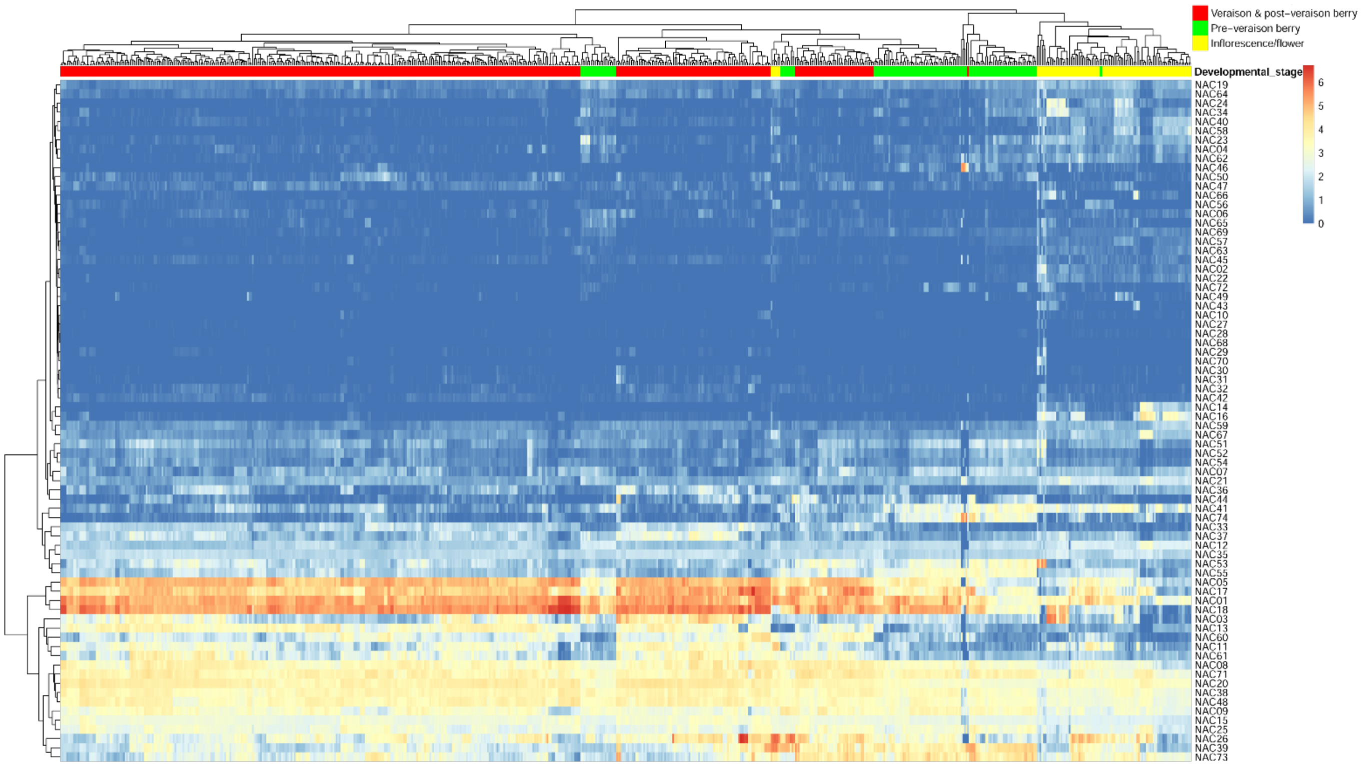

**Fig. S1.** Expression data were normalized by FPKM. Gene names are displayed to the right of each row. Samples were hierarchically clustered based on the Euclidean distance.

**Fig. S2**

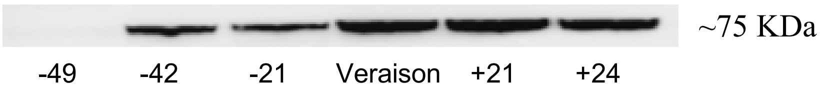

**Fig. S2.** Western blot analysis of different stages of berry development. Total protein extracts were blotted using anti-CARPO polyclonal antibody. Stages correspond to 49-, 42- and 21 days before veraison, veraison, 21- and 42 days post veraison. The bands (~75 KDa) represent the CARPO homodimer.

**Fig. S3**

```
                             NLS
                          ________
Pinot       MDNPQSTLPPGFRFHPTDEELILHYLSKKVTSTPFPVSIIADVDIYKFDPWELPGKAVFG    60
Syrah       MDNPQSTLPPGFRFHPTDEELILHYLSKKVTSTPFPVSIIADVDIYKFDPWELPGKAVFG    60
            ************************************************************

Pinot       EKEWYFFSPRDRKYPNGLRPNRAAASGYWKATGTDKTIVAASSIGGGGHGHIGVKKALVF    120
Syrah       EKEWYFFSPRDRKYPNGLRPNRAAASGYWKATGTDKTIVAASSIGGGGHGHIGVKKALVF    120
            ************************************************************

Pinot       YQGRPPKGIKTNWIMHEYRLAQPPNPAINKPPLKLRDASMRLDNWVLCRIYKKSNAVPPA    180
Syrah       YQGRPPKGIKTNWIMHEYRLAQPPNPAINKPPLKLRDASMRLDNWVLCRIYKKSNAVPPA    180
            ************************************************************

Pinot       TAAAIDDDREQEDSFMEESLKSHPNQSTIQPQKPSSFSNILDAMDSSTLGHLFSDIQYSD    240
Syrah       TAAAIDDDREQEDSFMEESLKSHPNQSTIQPQKPSSFSNILDVMDSSTLGHLFSDIQYSD    240
            ******************************************.*****************

Pinot       PTGFEPTPPAKYGSLGQSNDILPKLPYWKSVPSMENQLKRQRSSMDGDVPCPSKKLTSSC    300
Syrah       PTGFEPTPPAKYGSLSQSNDILPKLPYWKSVPSMENQLKRQRSSMDGDVPCPSKKLTSSC    300
            ***************.********************************************

Pinot       TFTTNPNQSDLPQSYFNQSLFNQALLLNPYFPFQG*     335
Syrah       TFTTNPNQSDLPQSYFNQSLFNQALLLNPYFPFQG*     335
            ***********************************
```

**Fig. S3.** Alignment of CARPO predicted amino acidic sequences from Pinot Noir and Syrah cultivars. Different amino acids are highlighted in yellow. The black line above the alignment locates the nuclear localization signal (NLS) predicted by PSORTII. The predicted aminoacidic residues located at the dimer interface are labeled in red (when common to at least half of ref. (32) analyzed sequences) and green (when common to at least one the ref. (32) analyzed sequences).

**Fig. S4**

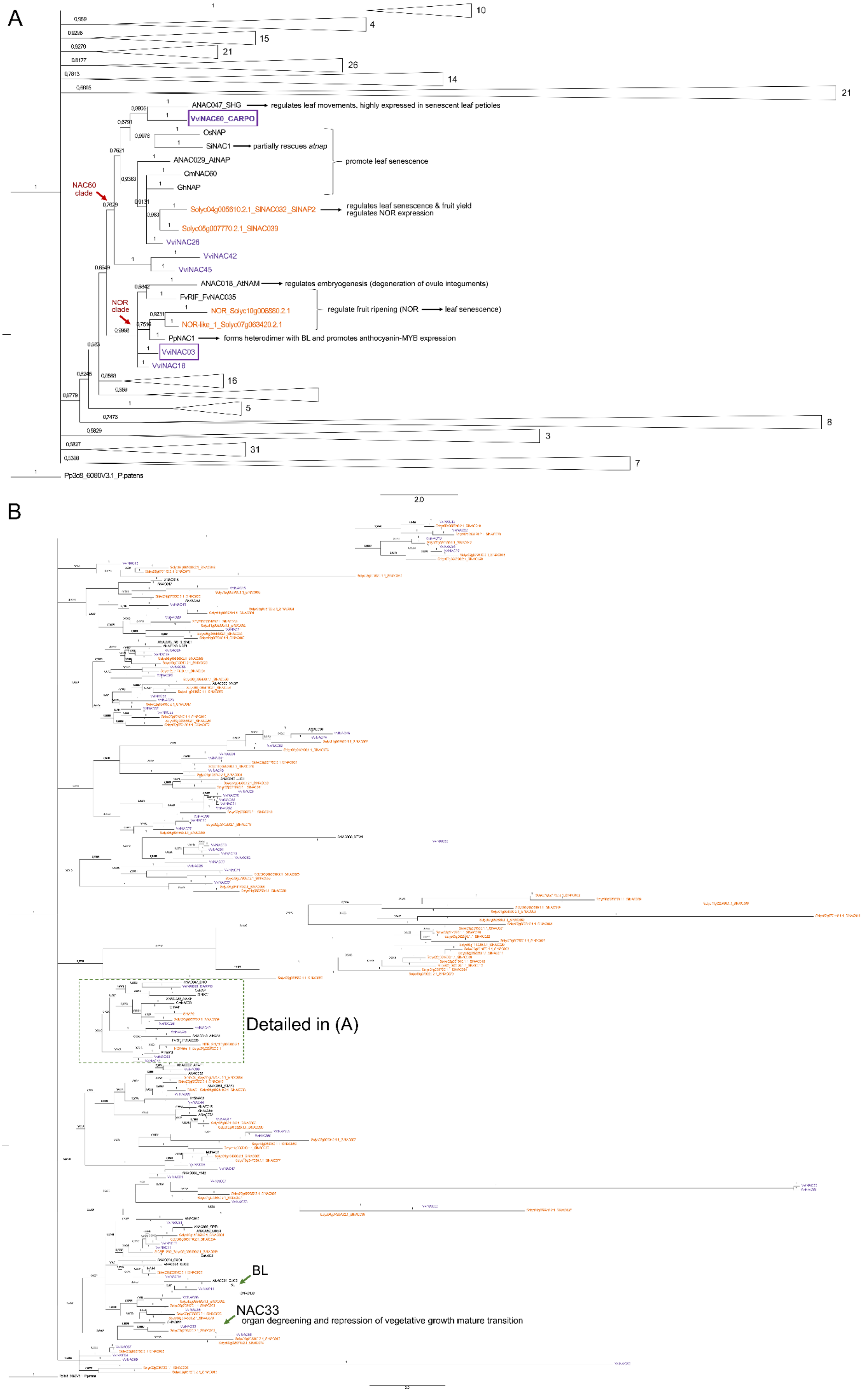
A
10
4
15
21
26
14
21
ANAC047_SHG
regulates leaf movements, highly expressed in senescent leaf petioles
VvNAC60_CARPO
OsNAP
SlNAC1
partially rescues atnap
ANAC029_AtNAP
CmNAC60
GhNAP
promote leaf senescence
NAC60 clade
Solyc04g005610.2.1_SlNAC032_SlNAP2
regulates leaf senescence & fruit yield
regulates NOR expression
Solyc05g007770.2.1_SlNAC039
VvNAC26
VvNAC42
VvNAC45
ANAC018_AtNAM
regulates embryogenesis (degeneration of ovule integuments)
FvRIF_FvNAC035
NOR clade
NOR_Solyc10g006880.2.1
NOR-like_1_Solyc07g063420.2.1
regulate fruit ripening (NOR
leaf senescence)
PpNAC1
forms heterodimer with BL and promotes anthocyanin-MYB expression
VvNAC03
VvNAC18
16
5
8
3
31
7
Pp3c8_6080V3.1_P.patens
2.0
B
Detailed in (A)
BL
NAC33
organ degreening and repression of vegetative growth mature transition

**Fig. S4.** Phylogenetic relationships of NAC gene of different plant species. Phylogenetic tree of NAC genes in grapevine, tomato and Arabidopsis and all characterized NACs in any other species. *V. vinifera* proteins are highlighted in purple colour, *S. lycopersicum* in orange and other species in black. Branch numbers depicts the posterior probability for clade. The big resolution figures is at: https://tomsbiolab.com/wp-content/uploads/2021/10/Fig.-S4.png.

**Fig. S5**

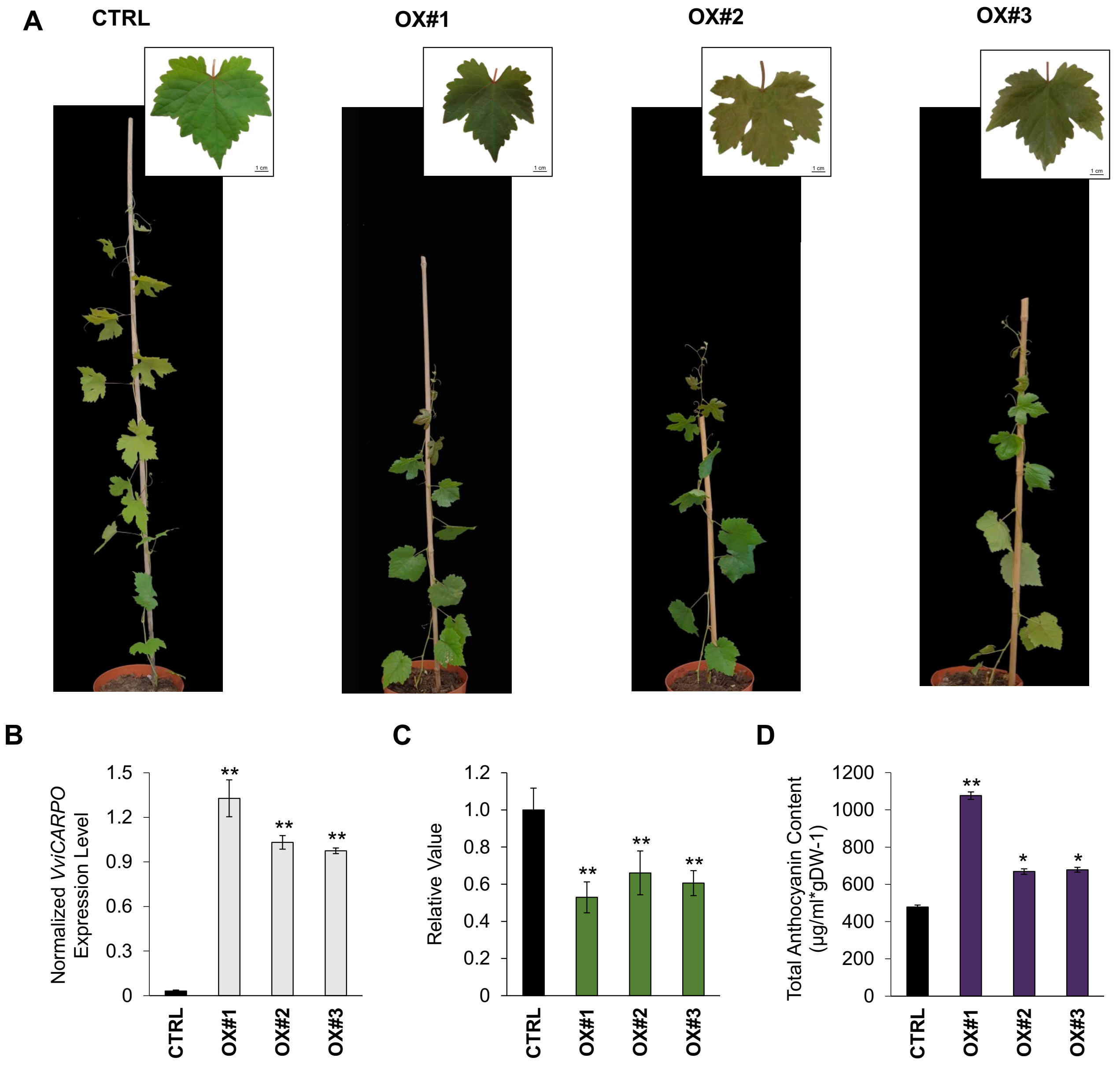

**Fig. S5.** Phenotypic changes in transgenic grapevine plants overexpressing *CARPO*. (*A*) Whole-plant and fully expanded leaf phenotypes caused by the ectopic expression of *CARPO* in the three independent OX.CARPO lines (OX#1, OX#2 and OX#3) compared to the vector control. We selected line #1 for further analysis (Fig. 2). (*B*) *CARPO* expression level by qPCR in OX.CARPO mature leaves. Each value corresponds to the mean ± SD of three technical replicates relative to the *VviUBIQUITIN1* (VIT_16s0098g01190) control. CTRL, control line; OX#1–#3, OX.CARPO transgenic lines. Asterisks indicate significant differences (**, P < 0.01; *t*-test) in the OX.CARPO lines compared to the control. SD, standard deviation. (*C*) Relative leaf area measurement of OX.CARPO lines compared to the control (n = 12 ± SD). Asterisks indicate significant differences (**, P < 0.01; *t*-test) in the OX.CARPO lines compared to the vector control. SD, standard deviation.

(*D*) Total anthocyanin content in leaves of OX.CARPO lines (n = 14 ± SD). Asterisks indicate significant differences (*, $P < 0.05$; **, $P < 0.01$; *t*-test) in the CARPO.EAR lines compared to the vector control. SD, standard deviation.

**Fig. S6**

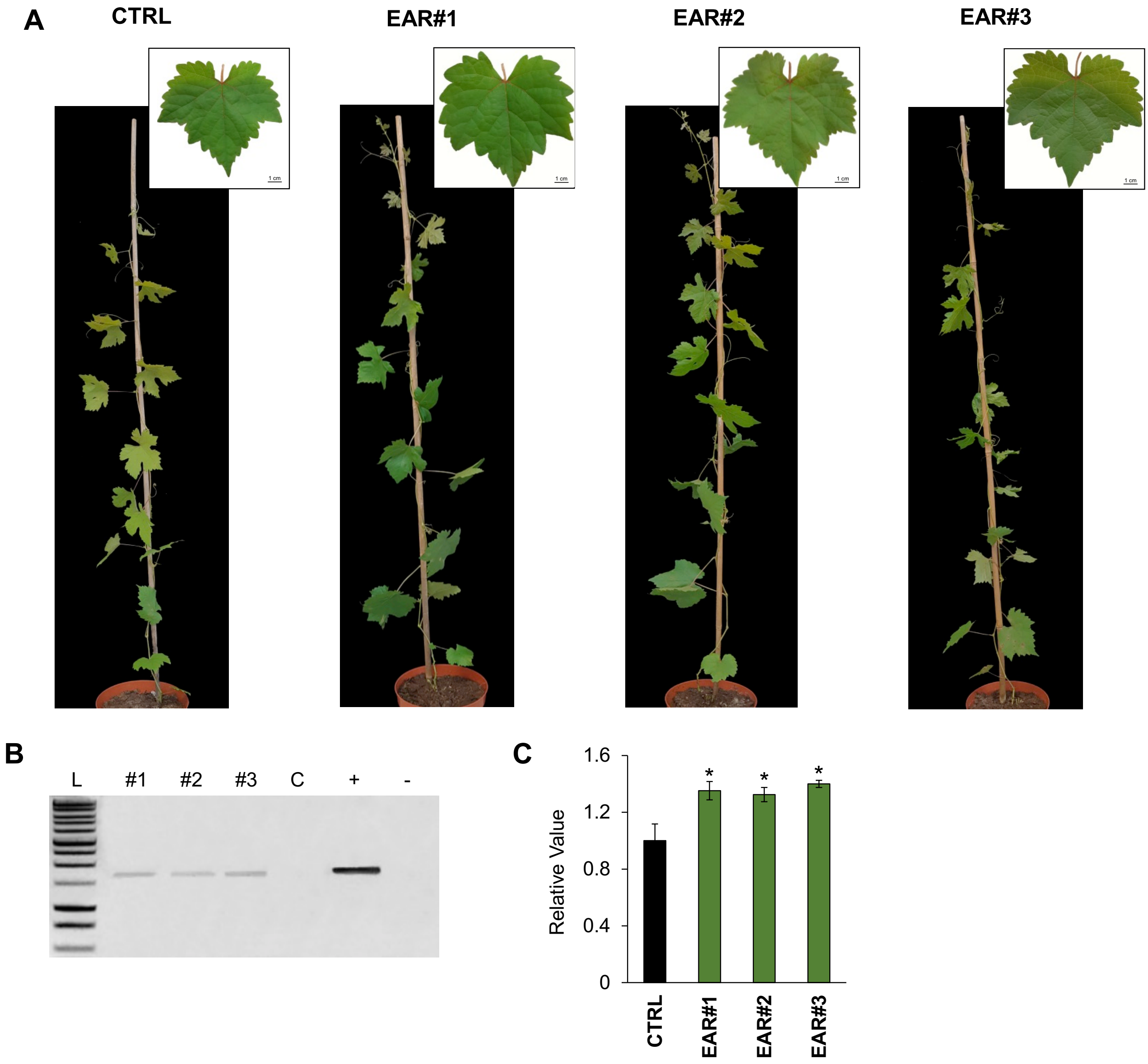

**Fig. S6.** Phenotypic changes in transgenic grapevine plants expressing the repressor CARPO-EAR. (*A*) Whole-plant and fully expanded leaf phenotypes caused by the ectopic expression of *CARPO-EAR* in the three independent CARPO.EAR lines (#1, #2 and #3) compared to the vector control. We selected line #3 for further analysis (Fig. 2). (*B*) Reverse transcription-PCR (RT-PCR) analysis of VviCARPO transcripts in fully expanded leaves of CARPO.EAR lines and the vector control. L, 1 kb ladder; #1-#3, CARPO.EAR lines; C, control line; +, positive control; -, negative control (the modified pK7WG2 plasmid; ***SI Materials and Methods***). We selected #3 (CARPO.EAR) for further analysis. (*C*) Relative leaf area measurement of CARPO.EAR lines compared to the control (n = 12 ± SD). Asterisks indicate significant differences (*, $P < 0.01$; *t*-test) in the CARPO.EAR lines compared to the vector control. SD, standard deviation.

**Fig. S7**

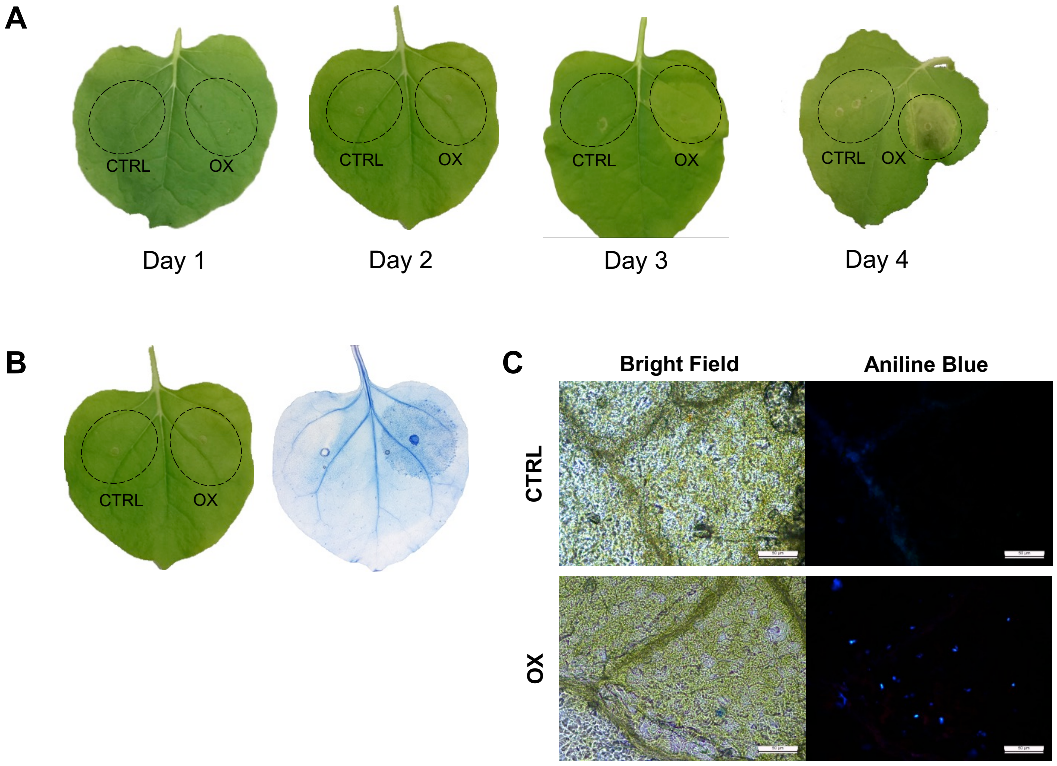

**Fig. S7.** Effects of transient overexpression of CARPO in *N. benthamiana* leaves. (*A*) Phenotype of *N. benthamiana* leaves agroinfiltrated with 35S:CARPO (OX) and the control (CTRL) on days 1, 2, 3 and 4 post-infiltration. (*B*) Phenotype showed by *N. benthamiana* leaf after 48 hours from the agroinfiltration with 35S:CARPO (OX) and the control (CTRL) (left) and cell death visualized with trypan blue staining (right). (C) Callose deposition on *N. benthamiana* leaves after 48 hours from the agroinfiltration with 35S:CARPO (OX) and the control (CTRL) visualized by aniline blue staining in a bright field (on the left) and under a fluorescence microscope (on the right). Magnification 20x.

**Fig. S8**

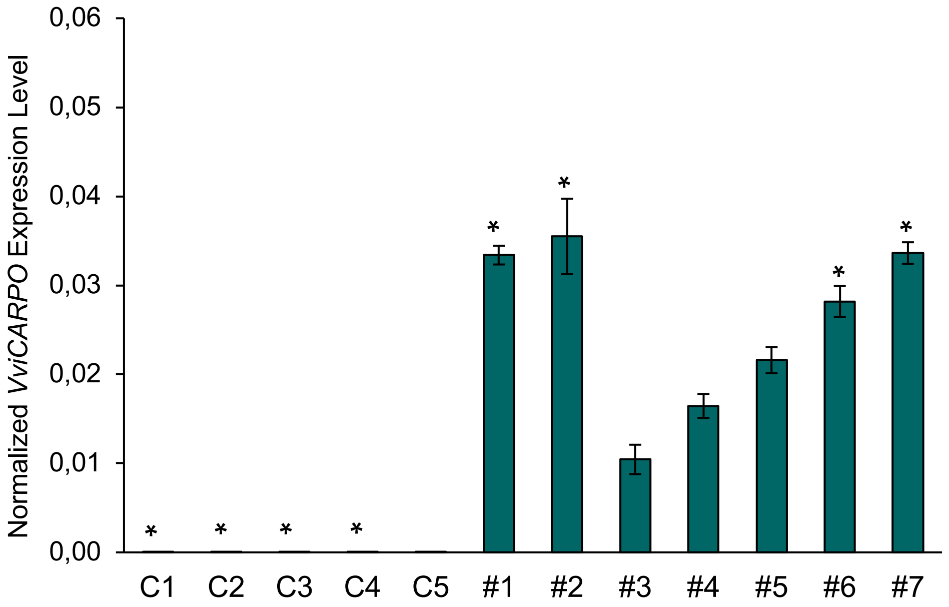

**Fig. S8.** *CARPO* expression level by qPCR in transgenic grapevine cv. Sultana leaves. Each value corresponds to the mean ± SD of three technical replicates relative to the *VviUBIQUITIN1* (VIT_16s0098g01190) control. C1-C5, control lines; #1-#7, overexpressing lines. Asterisks (*) indicate the selected lines for further analysis. SD, standard deviation.

**Fig. S9**

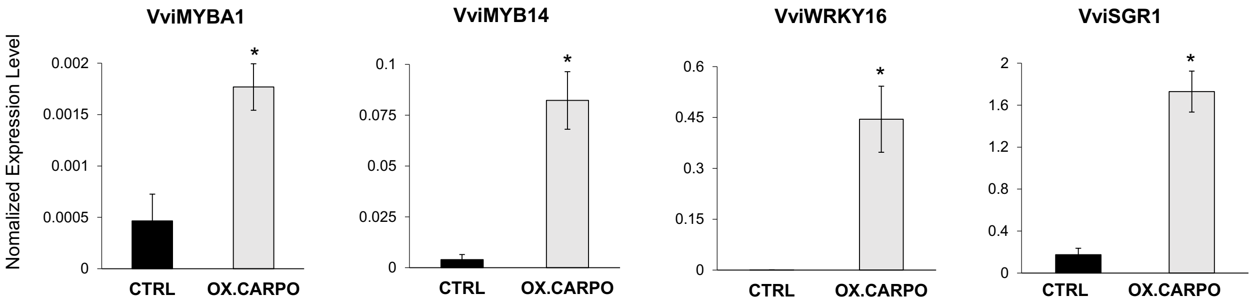

**Fig. S9.** Expression level of *VviMYBA1*, *VviMYB14*, *VviWRKY16* and *VviSGR1* by qPCR in transgenic *CARPO* overexpressing grapevine cv. Syrah leaves. Each value corresponds to the mean ± SD of three technical replicates relative to the *VviUBIQUITIN1* (VIT_16s0098g01190) control. CTRL, control lines; OX.CARPO, *CARPO* overexpressing line. Asterisks indicate significant differences (*, $P < 0.01$; *t*-test) in OX.CARPO leaves compared to the control. SD, standard deviation.

#### Fig. S10

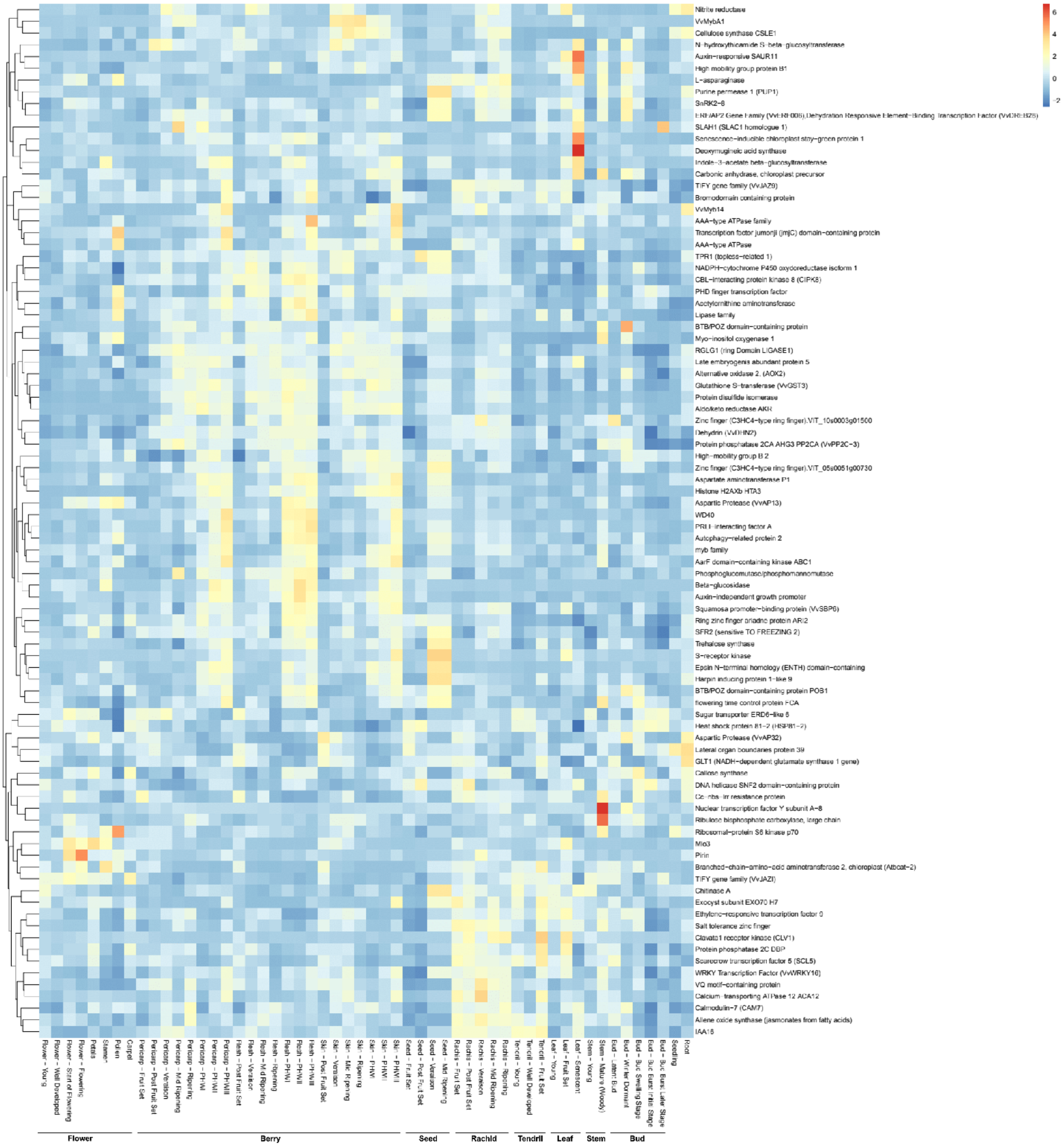

**Fig. S10.** Expression profiles of CARPO VHCT genes by exploring the cv. Corvina atlas dataset. The abbreviations after organ correspond to: FS, fruit set; PFS, post fruit set; V, véraison; MR, mid-ripening; R, ripening; Bud - L, latent bud; Bud - W, winter bud; Bud - S, bud swell; Bud - B, bud burst; Bud - AB, bud after burst; Inflorescence - Y, young; Inflorescence - WD, well-developed; Flower - FB, flowering begins; Flower - F, flowering; Tendril - Y, young; Tendril - WD, well-developed; Tendril - FS, mature; Leaf - Y, young; Leaf - FS, mature; Leaf - S, senescing leaf; Stem - G, green; Stem - W, woody.

**Fig. S11**

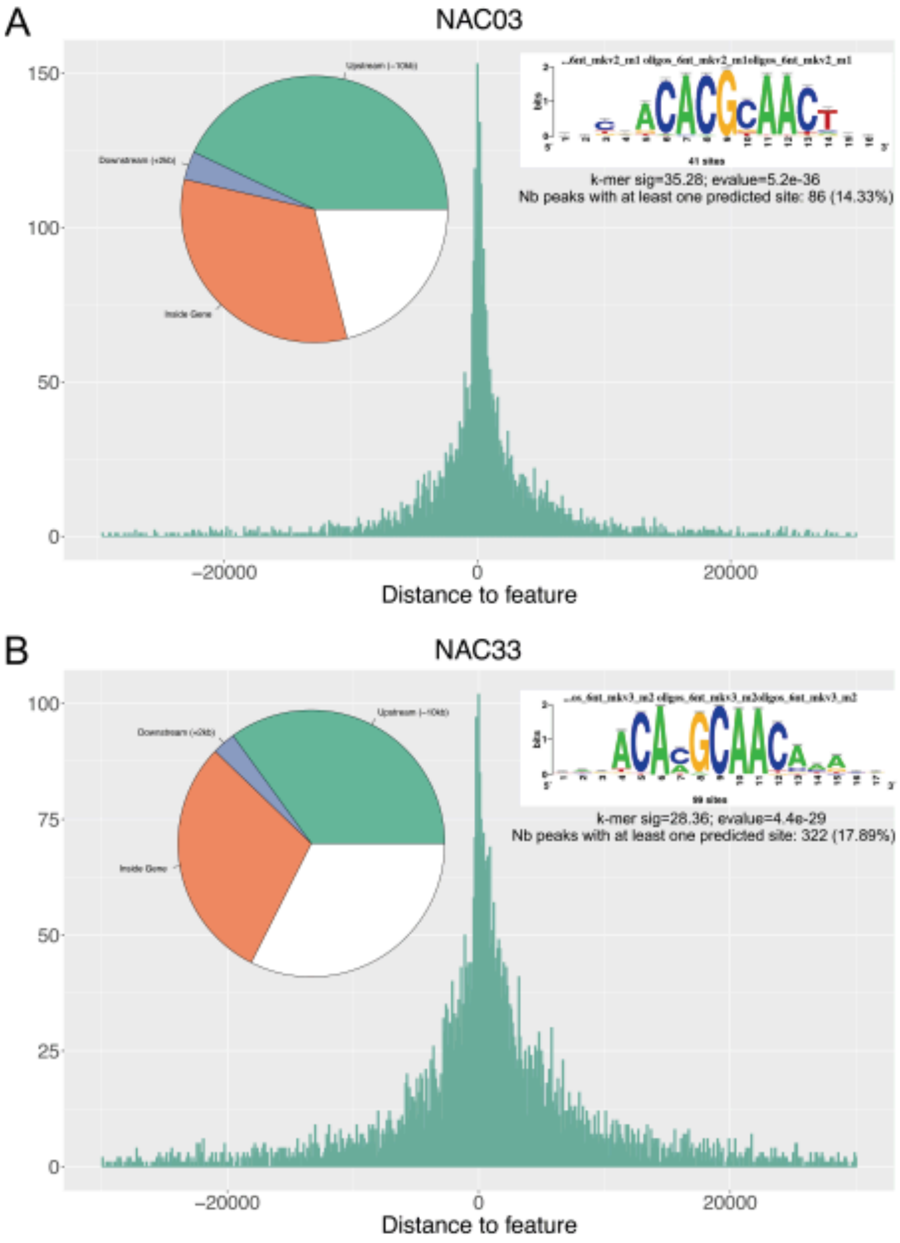

**Fig. S11.** Distribution of (A) NAC03 and (B) NAC33 DNA binding events with respect to their position from the transcription start sites (TSS) of their assigned genes in the leaf500 analysis. Distribution of peak positions is represented with a piechart for each NAC. De novo forward binding motif obtained from the inspection of the top 600-scoring peaks of each NAC TF were obtained using RSAT suite with default parameters as for CARPO.

**Fig. S12**

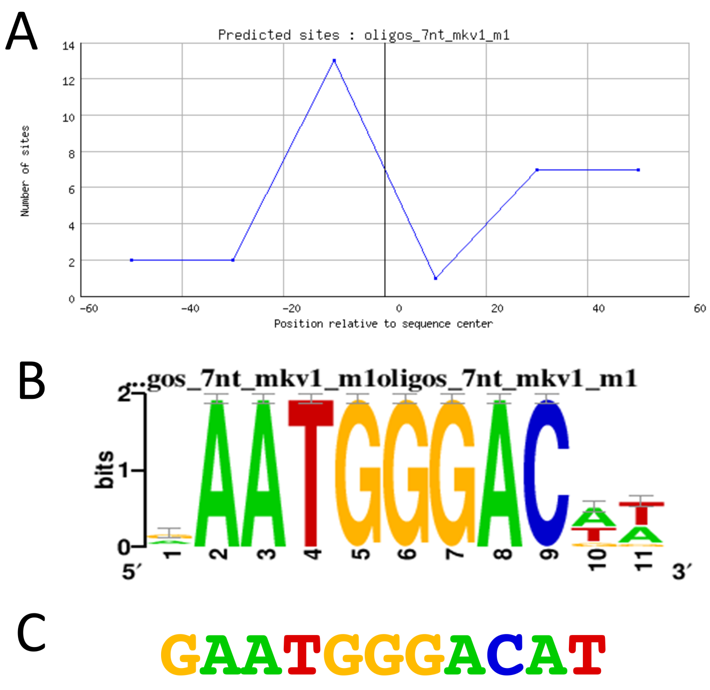

**Fig. S12**. Identification of a novel CARPO-binding motif in the proximal promoter regions of MYBA genes from chromosome 2 and 14. (A) Position of the motif with respect to the center of the binding regions (100bp) (B) Consensus de novo forward binding motif identified by RSAT, obtained from the inspection of CARPO-binding events in the proximal promoters of MYBA1/A2/A3/A5/A6/A7/A12 genes. (C) Motif sequence in the promoter of MYBA1 found in all three analyses (leaf500, leaf1000 and berry500).

**Fig. S13**

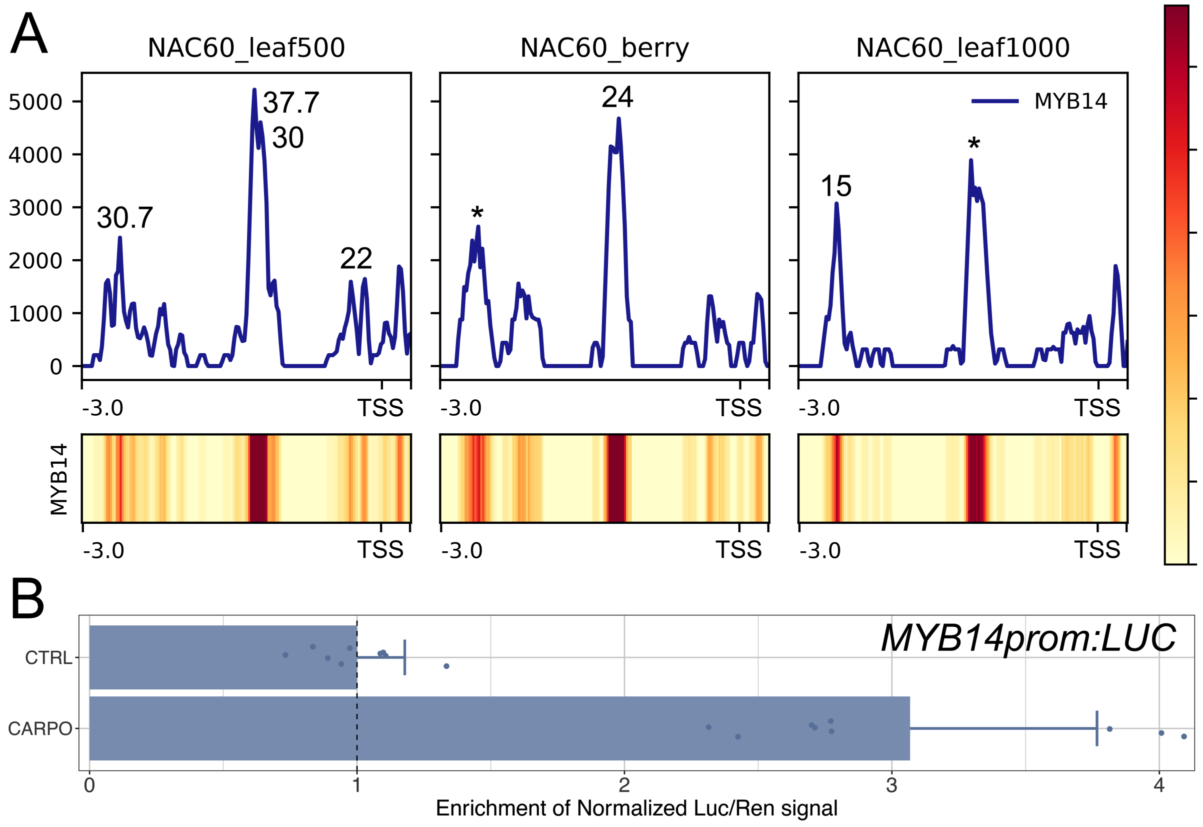

**Fig. S13**. (*A*) NAC DNA binding landscapes in the proximal promoter region of Myb14 gene. (*B*) Myb14 promoter activation tested by dual-luciferase reporter assay in infiltrated *N. benthamiana* leaves.

**Fig. S14**

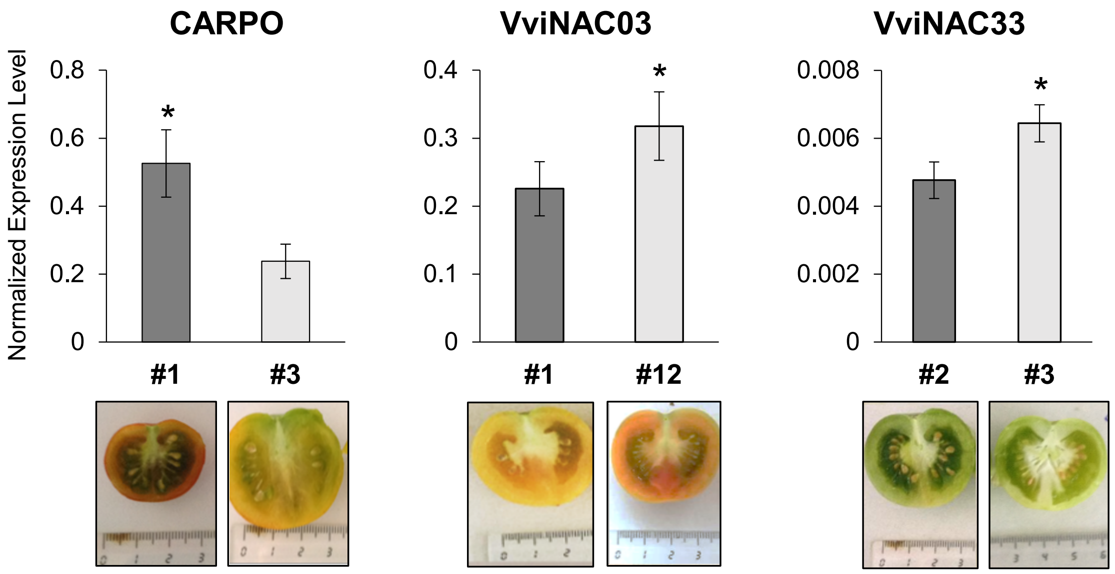

**Fig. S14.** Expression level of each transgene (CARPO, VviNAC03 and VviNAC33) by qPCR in $T_3$ fruits in *nor* mutant background at Br+7. Each value corresponds to the mean ± SD of three technical replicates relative to the ACTIN (Solyc03g078400) control. Phenotype of tomato fruits, corresponding to each line per transgene, are reported below each graph. Asterisks (*) indicate the selected lines for further analysis. SD, standard deviation.

**Fig. S15**

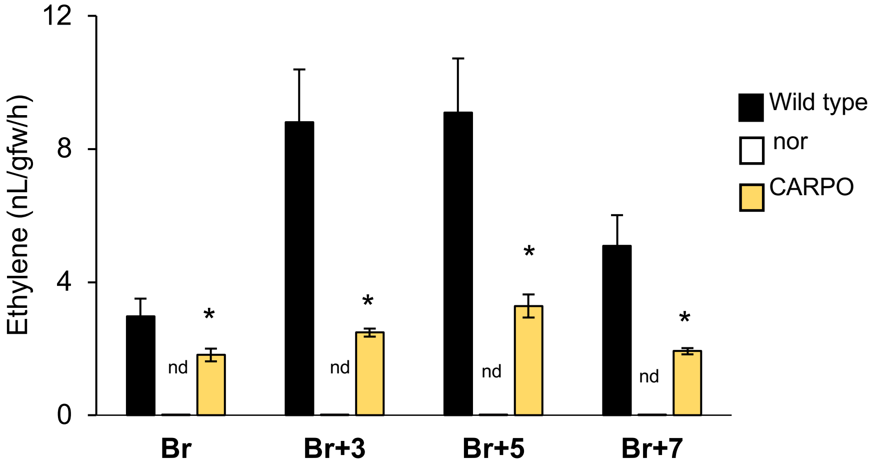

**Fig. S15.** Ethylene production during ripening of tomato in wild type, *nor* and $T_3$ fruit transformed with 35S:CARPO (CARPO) in *nor* tomato mutant background. Each value represents the mean ± SD of three biological replicates. Asterisks indicate significant differences (*, $P < 0.05$; *t*-test) in *CARPO* fruits compared to the *nor*. Nd, not detected; SD, standard deviation.

**Fig. S16**

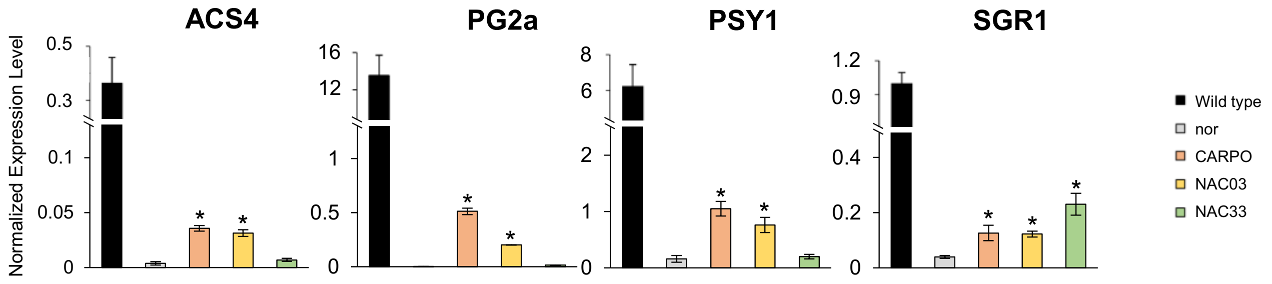

**Fig. S16.** Expression levels of tomato ripening-related genes (*ACS4; PG2a; PSY1; SGR1*) by qPCR in WT, *nor* and $T_3$ fruit transformed with 35S:*CARPO* (CARPO), 35S:*VviNAC03* (NAC03) and 35S:*VviNAC33* (NAC33) in *nor* tomato mutant background at Br+3. Each value corresponds to the mean ± SD of three technical replicates relative to the ACTIN (Solyc03g078400) control. Asterisks indicate significant differences (*, P < 0.01; *t*-test) in CARPO transgenic fruits compared to the *nor*. ACS4, 1-aminocyclopropane-1-carboxylic acid (ACC) synthase; PG2a, polygalacturonase 2A; PSY1, phytoene synthase 1; SGR1, stay-green protein 1; SD, standard deviation.

#### Fig. S17

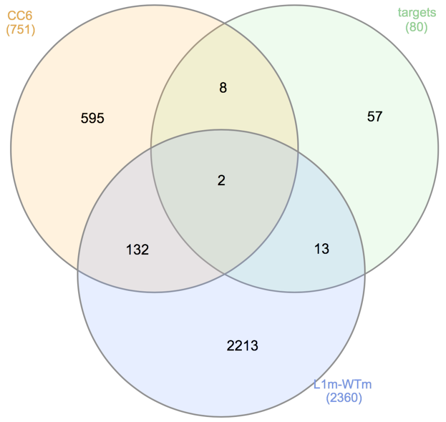

| [CC6] and [targets] and [L1m-WTm]: | Annotation |
|---|---|
| VIT_00s0179g00260 | Calcium-transporting ATPase 12 ACA12 |
| VIT_08s0007g02360 | Harpin inducing protein 1-like 9 |
| **[CC6] and [targets]:** | Annotation |
| VIT_18s0001g10150 | ERF/AP2 Gene Family (VvERF006),Dehydration Responsive Element-Binding Transcription Factor (VvDREB28) |
| VIT_12s0028g03270 | Ethylene-responsive transcription factor 9 |
| VIT_06s0004g01210 | PRLI-interacting factor A |
| VIT_18s0001g09230 | Salt tolerance zinc finger |
| VIT_11s0016g00710 | TIFY gene family (VvJAZ9) |
| VIT_05s0051g00730 | VviATL104 (defense) |
| VIT_07s0005g03340 | VvMyb14 |
| VIT_06s0004g07500 | WRKY Transcription Factor (VvWRKY16) |
| **[targets] and [L1m-WTm]:** | Annotation |
| VIT_17s0000g05420 | AAA-type ATPase family |
| VIT_18s0001g11630 | Allene oxide synthase (jasmonates from fatty acids) |
| VIT_04s0008g03770 | Aspartate aminotransferase P1 |
| VIT_06s0061g01030 | Aspartic Protease (VvAP13) |
| VIT_15s0048g00530 | Auxin-responsive SAUR11 |
| VIT_06s0004g04590 | Epsin N-terminal homology (ENTH) domain-containing protein |
| VIT_06s0004g07230 | Indole-3-acetate beta-glucosyltransferase |
| VIT_07s0005g00660 | Late embryogenis abundant protein 5 |
| VIT_18s0001g08510 | Lipase family |
| VIT_19s0090g01530 | Pirin |
| VIT_04s0044g01620 | RGLG1 (ring Domain LIGASE1) |
| VIT_13s0156g00590 | S-receptor kinase |
| VIT_10s0116g00960 | Transcription factor jumonji (jmjC) domain-containing protein |

**Fig. S17.** CARPO VHCT found in the module of ATL co-expressed genes specifically related to biotic stress (CC6, (33)) and/or up-regulated in grapevine plants overexpressing VviATL156 (L1mvsWTm; (34)).

**Fig. S18**

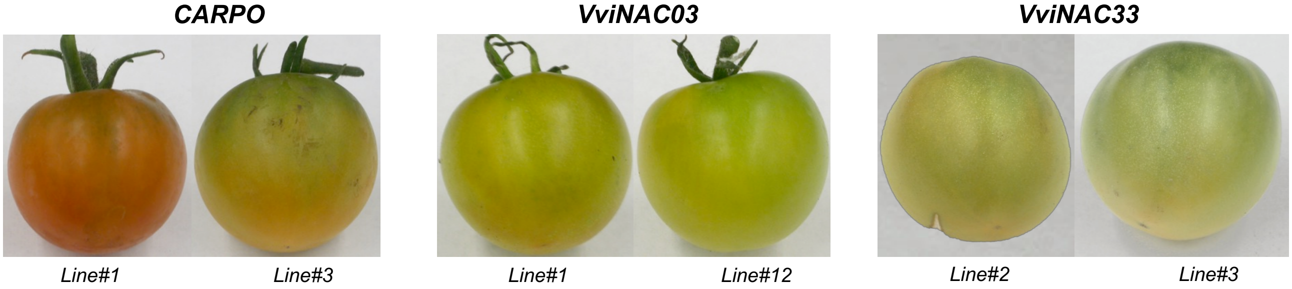

**Fig. S18.** Phenotype of the two selected lines of $T_0$ tomato fruits (*Solanum lycopersicum* cv. Ailsa Craig) in *nor* tomato mutant background for 35S:*CARPO* (CARPO), 35S:*VviNAC03* (VviNAC03) and 35S:*VviNAC33* (VviNAC33).

**Fig. S19**

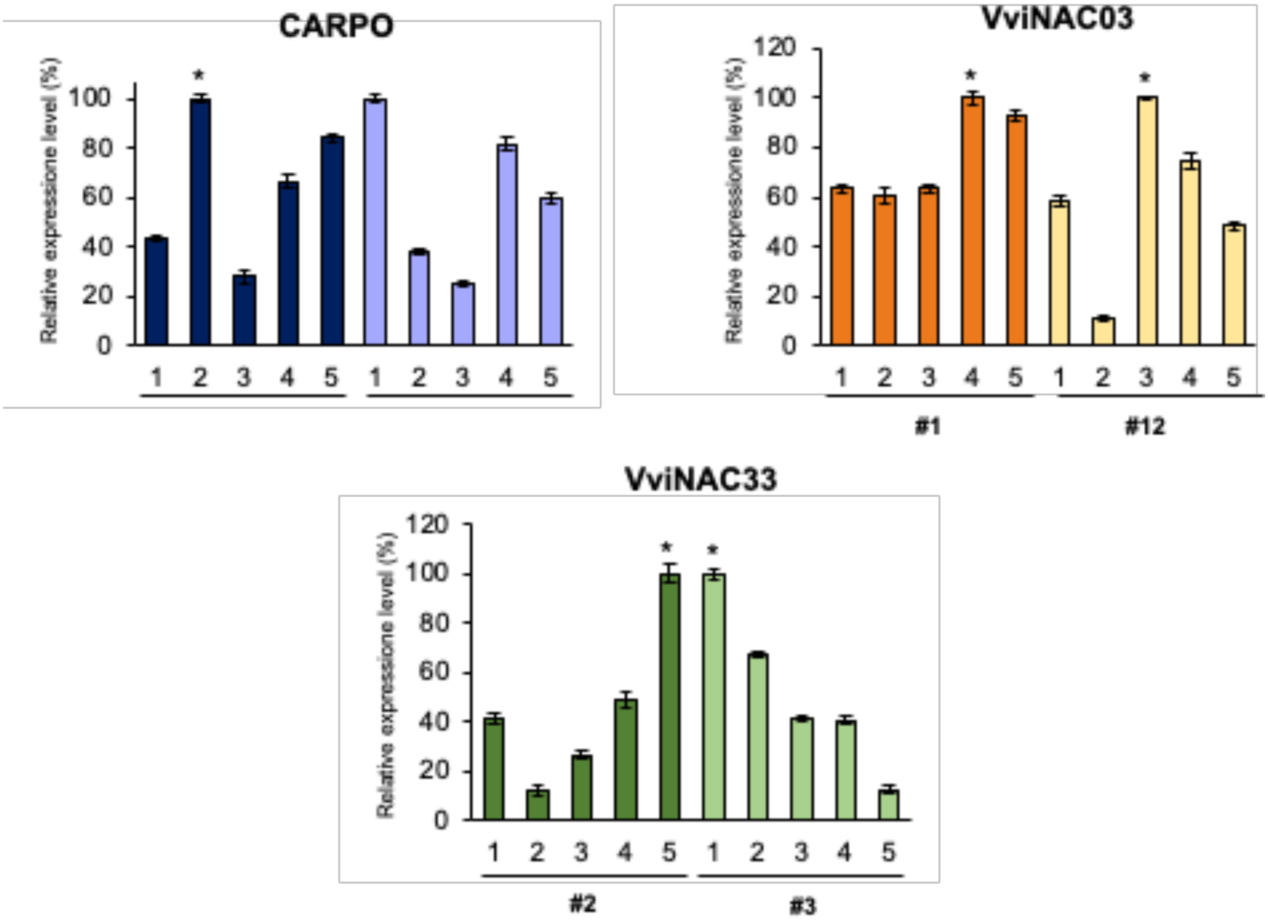

**Fig. S19.** Expression level of each transgene by qPCR in $T_1$ leaves in *nor* tomato mutant background. Each value corresponds to the mean ± SD of three technical replicates relative to the ACTIN (Solyc03g078400) control and normalized on the expression level of the maximum one of both lines for each transgene. Asterisks (*) indicate the selected lines for further analysis. SD, standard deviation.

#### Supplemental Tables

**Tab. S1.** The 89 very high confident targets (VHCTs) genes identified by combining DAP-seq with transcriptomics data.

| Gene ID | Distances for leaf 500 (r1+r2) | Distances for leaf 1000 | Distances for berry | FC Stable OX | FC Transient OX | Functional annotation |
|---|---|---|---|---|---|---|
| VIT_18s0001g08430 | -550 | -548 | -549 | 11.12 | | Branched-chain-amino-acid aminotransferase 2, chloroplast (Atbcat-2) |
| VIT_01s0011g00760 | -880 | -878 | -883 | 7.82 | | Beta-glucosidase |
| VIT_07s0005g00660 | -738 | -740 | | 6.76 | | Late embryogenis abundant protein 5 |
| VIT_06s0004g04590 | -92 | -90 | -91 | 6.49 | | Epsin N-terminal homology (ENTH) domain-containing |
| VIT_02s0025g04660 | -374 | -327 | -328 | | 5.74 | Senescence-inducible chloroplast stay-green protein 1 |
| VIT_07s0129g00330 | 37 | 35 | | 4.7 | | Lateral organ boundaries protein 39 |
| VIT_18s0001g13450 | -788 | -786 | -787 | 3.5 | | SLAH1 (SLAC1 homologue 1) |
| VIT_06s0004g07230 | -191 | -193 | -192 | | 3.45 | Indole-3-acetate beta-glucosyltransferase |
| VIT_03s0063g00370 | -685 | -683 | -684 | 3.38 | | Nitrite reductase |
| VIT_08s0040g02180 | -353 | -326 | | 3.28 | | Mlo3 |
| VIT_17s0000g07790 | -346 | -348 | | 3.22 | | N-hydroxythioamide S-beta-glucosyltransferase |
| VIT_17s0000g05420 | -37 ; -970 | -39 ; -967 | | 3.18 | | AAA-type ATPase family |
| VIT_11s0149g00280 | -810 | -799 | -823 | 3.12 | | Chitinase A |
| VIT_07s0005g03340 | -1377 | | -1358 | 2.88 | | VvMyb14 |
| VIT_01s0011g06460 | -46 ; 10 | -48 ; 12 | | 2.86 | | Deoxymugineic acid synthase |
| VIT_15s0048g00530 | -663 | -661 | | 2.85 | | Auxin-responsive SAUR11 |
| VIT_02s0033g00410 | -10 | -12 | | 2.83 | | VvMybA1 |
| VIT_13s0019g01980 | 40 | -12 | 16 | 2.77 | | Aspartic Protease (VvAP32) |
| VIT_01s0127g00590 | 44 | 44 | 43 | 2.75 | | Protein disulfide isomerase |
| VIT_00s0304g00030 | -938 | -940 | -939 | 2.69 | | VQ motif-containing protein |
| VIT_04s0008g00430 | -658 | -651 | | 2.69 | | Clavata1 receptor kinase (CLV1) |
| VIT_10s0003g01500 | -1016 | -1014 | -1015 | 2.63 | | Zinc finger (C3HC4-type ring finger) |
| VIT_11s0016g00710 | -101 | -99 | -100 | 2.55 | | TIFY gene family (VvJAZ9) |
| VIT_06s0004g04010 | -98 | -100 | -99 | 2.45 | | Exocyst subunit EXO70 H7 |
| VIT_19s0015g00710 | -381 | -381 | -394 | 2.34 | | Cellulose synthase CSLE1 |
| VIT_06s0004g07500 | -581 | -583 | | 2.27 | | WRKY Transcription Factor (VvWRKY16) |
| VIT_13s0156g00590 | -234 | -236 | | | 2.23 | S-receptor kinase |
| VIT_18s0001g09910 | -108 | -106 | | | 2.17 | L-asparaginase |
| VIT_09s0002g01590 | -459 | -458 | -457 | 2.15 | | Nuclear transcription factor Y subunit A-8 |
| VIT_04s0008g03770 | -143 | -141 | -248 | 2.14 | | Aspartate aminotransferase P1 |
| VIT_19s0090g01530 | -1008 | -1006 | | 2.14 | | Pirin |
| VIT_18s0001g06310 | -5 ; -46 | -48 ; -7 | -6 | 2.12 | | SnRK2-8 |
| VIT_18s0001g07400 | -318 | -320 | -319 | 2.11 | | Carbonic anhydrase, chloroplast precursor |
| VIT_04s0008g06330 | -155 | -114 | | 2.08 | | TPR1 (topless-related 1) |
| VIT_18s0001g11630 | -307 | -305 | | 2.06 | | Allene oxide synthase (jasmonates from fatty acids) |
| VIT_12s0028g03270 | 68; -279 | 66 ; -277 | 67 | 2.02 | | Ethylene-responsive transcription factor 9 |
| VIT_07s0005g02110 | -674 | -672 | | 1.99 | | Protein phosphatase 2C DBP |
| VIT_00s0179g00260 | -481 | -479 | | 1.96 | | Calcium-transporting ATPase 12 ACA12 |
| VIT_18s0001g09230 | -168 ; -1401 | -1399 ; -166 | | 1.95 | | Salt tolerance zinc finger |
| VIT_01s0011g05560 | -884 | -821 | -822 | 1.94 | | TIFY gene family (VvJAZI) |
| VIT_08s0007g02360 | -376 | -404 | | 1.92 | | Harpin inducing protein 1-like 9 |
| VIT_18s0001g10150 | -633 | -635 | | 1.9 | | ERF/AP2 Gene Family (VvERF006), Dehydration Responsive Element-Binding Transcription Factor (VvDREB28) |
| VIT_16s0098g00290 | -645 | -647 | -646 | 1.89 | | GLT1 (NADH-dependent glutamate synthase 1 gene) |
| VIT_04s0044g01620 | -171 | -134 | | 1.88 | | RGLG1 (ring Domain LIGASE1) |
| VIT_10s0003g05690 | -216 ; -1106 | -1104;-218 | -1097 ; -256 | | 1.88 | Ribulose bisphosphate carboxylase, large chain |
| VIT_18s0001g00360 | -324 ; -571 | -352 ; -569 | -570 ; -322 | 1.86 | | Dehydrin (VvDHN2) |
| VIT_04s0044g00110 | -394 | -396 | | 1.86 | | High-mobility group B 2 |
| VIT_02s0025g00880 | -736 | -734 | -735 | 1.83 | | BTB/POZ domain-containing protein POB1 |
| VIT_12s0028g00930 | 36 | 51 | - | 1.81 | | Glutathione S-transferase (VvGST3) |
| VIT_05s0062g00980 | 1 | -1 | -5 | 1.8 | | Aldo/keto reductase AKR |
| VIT_18s0122g01340 | -592 | -594 | - | 1.77 | | BTB/POZ domain-containing protein |
| VIT_05s0051g00730 | -51 | -49 | -131 | 1.75 | | Zinc finger (C3HC4-type ring finger) |
| VIT_18s0001g07630 | -233 | -407 | - | 1.75 | | NADPH-cytochrome P450 oxydoreductase isoform 1 |
| VIT_08s0007g02200 | -199 | -201 | - | 1.7 | | High mobility group protein B1 |
| VIT_11s0016g04490 | -1100 | -1104 | | 1.66 | | IAA16 |
| VIT_13s0019g02200 | -577 ; -797 | -579 ; -724 | - | 1.65 | | Protein phosphatase 2CA AHG3 PP2CA (VvPP2C-3) |
| VIT_13s0156g00210 | -13 | -15 | - | 1.65 | | Callose synthase |
| VIT_05s0077g00710 | 34 | 36 | - | 1.62 | | flowering time control protein FCA |
| VIT_09s0054g01620 | -186 | -184 | - | 1.59 | | myb family |
| VIT_18s0001g07980 | -423 | -383 | - | 1.59 | | CBL-interacting protein kinase 8 (CIPK8) |
| VIT_09s0002g02940 | -205 | -203 | -290 | 1.59 | | Myo-inositol oxygenase 1 |
| VIT_14s0083g00440 | -478 | -480 | - | 1.59 | | PHD finger transcription factor |
| VIT_00s0203g00100 | -302 | -300 | -301 | 1.57 | | AarF domain-containing kinase ABC1 |
| VIT_18s0001g11740 | -325 ; -401 | -399 ; -323 | | | 1.57 | Ring zinc finger ariadne protein ARI2 |
| VIT_09s0002g00270 | -718 | -716 | -717 | 1.56 | | R protein disease resistance protein |
| VIT_07s0104g00860 | -395 | -397 | -396 | 1.55 | | WD40 |
| VIT_10s0116g00960 | -909 ; -200 | -911 ; -198 | -198 | 1.55 | | Transcription factor jumonji (jmjC) domain-containing protein |
| VIT_06s0009g01630 | -478 | -480 | -479 | 1.52 | | Cc-nbs-lrr resistance protein |
| VIT_09s0002g08670 | -267 | -265 | -266 | 1.52 | | Acetylornithine aminotransferase |
| VIT_06s0004g01210 | -203 | -284 | | 1.49 | | PRLI-interacting factor A |
| VIT_17s0000g07520 | -1043 | -1052 | -1055 | 1.49 | | Ribosomal-protein S6 kinase p70 |
| VIT_15s0048g02990 | -617 | -615 | | 1.47 | | AAA-type ATPase |
| VIT_18s0001g06950 | -555 | -553 | | 1.45 | | Purine permease 1 (PUP1) |
| VIT_19s0090g01740 | -720 | -691 | | 1.45 | | Scarecrow transcription factor 5 (SCL5) |
| VIT_18s0001g08510 | -358 | -360 | -359 | 1.42 | | Lipase family |
| VIT_19s0176g00150 | 74 | 76 | 65 | 1.42 | | Phosphoglucomutase/phosphomannomutase |
| VIT_03s0038g03240 | -458 | -466 | | 1.4 | | Auxin-independent growth promoter |
| VIT_10s0003g01680 | -727 | -714 | -650 | 1.39 | | Trehalose synthase |
| VIT_15s0021g02180 | -56 | | -68 | | 1.39 | DNA helicase SNF2 domain-containing protein |
| VIT_03s0097g00570 | -137 | -135 | | | 1.38 | Bromodomain containing protein |
| VIT_12s0142g00430 | -344 | -346 | -345 | 1.38 | | Alternative oxidase 2, (AOX2) |
| VIT_05s0020g02160 | -175 | -168 | -135 | 1.37 | | Squamosa promoter-binding protein (VvSBP6) |
| VIT_18s0001g12560 | -186 | -145 | | 1.34 | | Sugar transporter ERD6-like 6 |
| VIT_05s0077g00550 | 33 | 31 | | 1.34 | | SFR2 (sensitive TO FREEZING 2) |
| VIT_06s0061g01030 | -247 | -245 | | 1.32 | | Aspartic Protease (VvAP13) |
| VIT_08s0040g00470 | 8 | 6 | | 1.32 | | Calmodulin-7 (CAM7) |
| VIT_07s0104g00970 | -176 | -200 | | 1.31 | | Histone H2AXb HTA3 |
| VIT_03s0063g00600 | -223 | -221 | | | 1.3 | Autophagy-related protein 2 |
| VIT_19s0015g01090 | -353 | -351 | | | 1.3 | Heat shock protein 81-2 (HSP81-2) |

### Tab. S2. Complete list of Gene Ontology terms of Figure 3D.

| query | significant | p_value | term_size | query_size | intersection_size | precision | recall | term_id | source | term_name | effective_domain_size | source_order | parents | intersection | logpval | opacity | term_size_scaled | order | .group |
|---|---|---|---|---|---|---|---|---|---|---|---|---|---|---|---|---|---|---|---|
| CARPO All peaks | TRUE | 1.72095380466079E-06 | 1705 | 6892 | 798 | 0.115786419 | 0.468035191 | GO:2001141 | GO:BP | regulation of RNA biosynthetic process | 17259 | 30327 | c(GO:0010556, GO:0031326, GO:0032774, GO:0051252) | VIT_05s0051g00450,VIT_00s0586g00010,VIT_08s0105g00230,VIT_18s0001g05890,VIT_17s0000g01280,VIT_15s0048g01740,VIT_02s0025g04460,VIT_14s0083g00700,VIT_04s0023g01490,VIT_01s0011g00100 | 5.764230787 | 0.8 | 7.502013216 | 193.3479593 | 1 |
| CARPO All peaks | TRUE | 1.72095380466079E-06 | 1705 | 6892 | 798 | 0.115786419 | 0.468035191 | GO:1903506 | GO:BP | regulation of nucleic acid-templated transcription | 17259 | 26558 | c(GO:0097659, GO:2001141) | VIT_05s0051g00450,VIT_00s0586g00010,VIT_08s0105g00230,VIT_18s0001g05890,VIT_17s0000g01280,VIT_15s0048g01740,VIT_02s0025g04460,VIT_14s0083g00700,VIT_04s0023g01490,VIT_01s0011g00100 | 5.764230787 | 0.8 | 7.502013216 | 178.6428071 | 1 |
| CARPO All peaks | TRUE | 1.79896911765533E-06 | 1692 | 6892 | 791 | 0.114770749 | 0.46749409 | GO:0006355 | GO:BP | regulation of transcription, DNA-templated | 17259 | 2372 | c(GO:0006351, GO:0010468, GO:1903506, GO:2000112) | VIT_05s0051g00450,VIT_00s0586g00010,VIT_08s0105g00230,VIT_18s0001g05890,VIT_17s0000g01280,VIT_15s0048g01740,VIT_02s0025g04460,VIT_14s0083g00700,VIT_04s0023g01490,VIT_01s0011g00100 | 5.744976292 | 0.8 | 7.496354062 | 84.2785694 | 1 |
| CARPO All peaks | TRUE | 8.57306938854857E-06 | 1795 | 6892 | 829 | 0.120284388 | 0.46183844 | GO:0051252 | GO:BP | regulation of RNA metabolic process | 17259 | 15773 | c(GO:0016070, GO:0019219, GO:0060255) | VIT_05s0051g00450,VIT_00s0586g00010,VIT_08s0105g00230,VIT_18s0001g05890,VIT_17s0000g01280,VIT_15s0048g01740,VIT_02s0025g04460,VIT_14s0083g00700,VIT_04s0023g01490,VIT_01s0011g00100 | 5.066863661 | 0.8 | 7.540047202 | 136.5639886 | 1 |
| CARPO All peaks | TRUE | 2.25345242365077E-05 | 4002 | 6892 | 1746 | 0.253337203 | 0.436281859 | GO:0050789 | GO:BP | regulation of biological process | 17259 | 15387 | c(GO:0008150, GO:0065007) | VIT_09s0054g00350,VIT_05s0051g00450,VIT_00s0586g00010,VIT_18s0001g01320,VIT_09s0096g00990,VIT_00s0396g00050,VIT_17s0000g02210,VIT_08s0105g00230,VIT_14s0060g02140,VIT_05s0062g01120 | 4.647151607 | 0.8 | 8.132879471 | 135.0579688 | 1 |
| CARPO All peaks | TRUE | 2.25345242365077E-05 | 4597 | 6892 | 1991 | 0.288885665 | 0.433108549 | GO:0065007 | GO:BP | biological regulation | 17259 | 18687 | GO:0008150 | VIT_09s0054g00350,VIT_05s0051g00450,VIT_00s0586g00010,VIT_18s0001g01320,VIT_09s0096g00990,VIT_00s0396g00050,VIT_17s0000g02210,VIT_08s0105g00230,VIT_14s0060g02140,VIT_05s0062g01120 | 4.647151607 | 0.8 | 8.235365623 | 147.9332679 | 1 |
| CARPO All peaks | TRUE | 2.25345242365077E-05 | 1897 | 6892 | 867 | 0.125798027 | 0.457037428 | GO:0010556 | GO:BP | regulation of macromolecule biosynthetic process | 17259 | 4728 | c(GO:0009059, GO:0009889, GO:0060255) | VIT_05s0051g00450,VIT_00s0586g00010,VIT_00s0396g00050,VIT_08s0105g00230,VIT_18s0001g05890,VIT_17s0000g01280,VIT_15s0048g01740,VIT_02s0025g04460,VIT_14s0083g00700,VIT_04s0023g01490 | 4.647151607 | 0.8 | 7.58091212 | 93.47075265 | 1 |
| CARPO All peaks | TRUE | 2.39882428224944E-05 | 1927 | 6892 | 879 | 0.127539176 | 0.456149455 | GO:0009889 | GO:BP | regulation of biosynthetic process | 17259 | 4230 | c(GO:0009058, GO:0019222) | VIT_05s0051g00450,VIT_00s0586g00010,VIT_00s0396g00050,VIT_08s0105g00230,VIT_18s0001g05890,VIT_17s0000g01280,VIT_15s0048g01740,VIT_02s0025g04460,VIT_14s0083g00700,VIT_04s0023g01490 | 4.620001564 | 0.8 | 7.592513613 | 91.52775296 | 1 |
| CARPO All peaks | TRUE | 2.47205710291047E-05 | 2108 | 6892 | 955 | 0.138566454 | 0.453036053 | GO:0010468 | GO:BP | regulation of gene expression | 17259 | 4670 | c(GO:0010467, GO:0060255) | VIT_05s0051g00450,VIT_00s0586g00010,VIT_00s0396g00050,VIT_08s0105g00230,VIT_18s0001g05890,VIT_17s0000g01280,VIT_15s0048g01740,VIT_02s0025g04460,VIT_14s0083g00700,VIT_04s0023g01490 | 4.606941502 | 0.8 | 7.658892231 | 93.24445951 | 1 |
| CARPO All peaks | TRUE | 2.56464495429008E-05 | 1880 | 6892 | 858 | 0.124492165 | 0.456382979 | GO:2000112 | GO:BP | regulation of cellular macromolecule biosynthetic process | 17259 | 29347 | c(GO:0010556, GO:0031326, GO:0034645) | VIT_05s0051g00450,VIT_00s0586g00010,VIT_00s0396g00050,VIT_08s0105g00230,VIT_18s0001g05890,VIT_17s0000g01280,VIT_15s0048g01740,VIT_02s0025g04460,VIT_14s0083g00700,VIT_04s0023g01490 | 4.590972749 | 0.8 | 7.574256224 | 189.5243856 | 1 |
| CARPO All peaks | TRUE | 2.57930143131912E-05 | 1921 | 6892 | 875 | 0.126958793 | 0.455491931 | GO:0031326 | GO:BP | regulation of cellular biosynthetic process | 17259 | 8296 | c(GO:0009889, GO:0031323, GO:0044249) | VIT_05s0051g00450,VIT_00s0586g00010,VIT_00s0396g00050,VIT_08s0105g00230,VIT_18s0001g05890,VIT_17s0000g01280,VIT_15s0048g01740,VIT_02s0025g04460,VIT_14s0083g00700,VIT_04s0023g01490 | 4.588497901 | 0.8 | 7.590207832 | 107.3916821 | 1 |
| CARPO All peaks | TRUE | 3.02160256170873E-05 | 1846 | 6892 | 842 | 0.122170633 | 0.456121343 | GO:0019219 | GO:BP | regulation of nucleobase-containing compound metabolic process | 17259 | 6505 | c(GO:0006139, GO:0031323, GO:0051171, GO:0080090) | VIT_05s0051g00450,VIT_00s0586g00010,VIT_08s0105g00230,VIT_18s0001g05890,VIT_17s0000g01280,VIT_15s0048g01740,VIT_02s0025g04460,VIT_14s0083g00700,VIT_04s0023g01490,VIT_01s0011g00100 | 4.51976266 | 0.8 | 7.560761943 | 100.4039061 | 1 |
| CARPO All peaks | TRUE | 3.02160256170873E-05 | 1855 | 6892 | 846 | 0.122751016 | 0.45606469 | GO:0097659 | GO:BP | nucleic acid-templated transcription | 17259 | 22023 | GO:0032774 | VIT_05s0051g00450,VIT_00s0586g00010,VIT_04s0079g00620,VIT_08s0105g00230,VIT_18s0001g05890,VIT_17s0000g01280,VIT_15s0048g01740,VIT_09s0002g07110,VIT_02s0025g04460,VIT_14s0083g00700 | 4.51976266 | 0.8 | 7.564357995 | 160.9490248 | 1 |
| CARPO All peaks | TRUE | 3.21626462696612E-05 | 1859 | 6892 | 847 | 0.122896111 | 0.455621302 | GO:0032774 | GO:BP | RNA biosynthetic process | 17259 | 9058 | c(GO:0009059, GO:0016070, GO:0034654) | VIT_05s0051g00450,VIT_00s0586g00010,VIT_04s0079g00620,VIT_08s0105g00230,VIT_18s0001g05890,VIT_17s0000g01280,VIT_15s0048g01740,VIT_09s0002g07110,VIT_02s0025g04460,VIT_14s0083g00700 | 4.492648226 | 0.8 | 7.565950643 | 110.3647057 | 1 |
| CARPO All peaks | TRUE | 3.98675123545306E-05 | 1845 | 6892 | 840 | 0.121880441 | 0.455284553 | GO:0006351 | GO:BP | transcription, DNA-templated | 17259 | 2368 | c(GO:0010467, GO:0034645, GO:0097659) | VIT_05s0051g00450,VIT_00s0586g00010,VIT_04s0079g00620,VIT_08s0105g00230,VIT_18s0001g05890,VIT_17s0000g01280,VIT_15s0048g01740,VIT_09s0002g07110,VIT_02s0025g04460,VIT_14s0083g00700 | 4.399380863 | 0.8 | 7.5603613 | 84.26296298 | 1 |
| CARPO All peaks | TRUE | 5.07168480270489E-05 | 3667 | 6892 | 1599 | 0.232008125 | 0.436051268 | GO:0050794 | GO:BP | regulation of cellular process | 17259 | 15391 | c(GO:0009987, GO:0050789) | VIT_09s0054g00350,VIT_05s0051g00450,VIT_00s0586g00010,VIT_18s0001g01320,VIT_00s0396g00050,VIT_17s0000g02210,VIT_08s0105g00230,VIT_05s0062g01120,VIT_18s0001g05890,VIT_17s0000g01280 | 4.294847745 | 0.8 | 8.06824203 | 135.0735752 | 1 |
| CARPO All peaks | TRUE | 0.000143618 | 2181 | 6892 | 976 | 0.141613465 | 0.447501146 | GO:0034654 | GO:BP | nucleobase-containing compound biosynthetic process | 17259 | 10049 | c(GO:0006139, GO:0018130, GO:0019438, GO:0044271, GO:19013 | VIT_05s0051g00450,VIT_00s0586g00010,VIT_04s0079g00620,VIT_00s0733g00010,VIT_08s0105g00230,VIT_11s0037g00950,VIT_18s0001g05890,VIT_17s0000g01280,VIT_15s0048g01740,VIT_09s0002g07110 | 3.842789819 | 0.8 | 7.684063805 | 114.2311971 | 1 |
| CARPO All peaks | TRUE | 0.00038097 | 2264 | 6892 | 1006 | 0.145966338 | 0.44434629 | GO:0051171 | GO:BP | regulation of nitrogen compound metabolic process | 17259 | 15706 | c(GO:0006807, GO:0019222) | VIT_05s0051g00450,VIT_00s0586g00010,VIT_09s0096g00990,VIT_00s0396g00050,VIT_08s0105g00230,VIT_18s0001g05890,VIT_17s0000g01280,VIT_15s0048g01740,VIT_02s0025g04460,VIT_14s0083g00700 | 3.419109654 | 0.8 | 7.711679656 | 136.302581 | 1 |
| CARPO All peaks | TRUE | 0.00038097 | 642 | 6892 | 314 | 0.04556007 | 0.489096573 | GO:0009725 | GO:BP | response to hormone | 17259 | 4093 | c(GO:0009719, GO:0010033) | VIT_08s0032g00440,VIT_00s0733g00010,VIT_05s0062g01120,VIT_02s0154g00010,VIT_05s0020g03310,VIT_03s0038g04390,VIT_17s0000g01830,VIT_14s0030g00440,VIT_17s0000g00320,VIT_08s0105g00570 | 3.419109654 | 0.8 | 6.779830517 | 90.99323297 | 1 |
| CARPO All peaks | TRUE | 0.000481347 | 646 | 6892 | 315 | 0.045705165 | 0.487616099 | GO:0009719 | GO:BP | response to endogenous stimulus | 17259 | 4087 | GO:0050896 | VIT_08s0032g00440,VIT_00s0733g00010,VIT_05s0062g01120,VIT_02s0154g00010,VIT_05s0020g03310,VIT_03s0038g04390,VIT_17s0000g01830,VIT_14s0030g00440,VIT_17s0000g00320,VIT_08s0105g00570 | 3.317542057 | 0.8 | 6.784422996 | 90.96982333 | 1 |
| CARPO All peaks | TRUE | 0.000512543 | 2321 | 6892 | 1028 | 0.149158445 | 0.442912538 | GO:0018130 | GO:BP | heterocycle biosynthetic process | 17259 | 6045 | c(GO:0044249, GO:0046483) | VIT_05s0051g00450,VIT_00s0586g00010,VIT_04s0079g00620,VIT_00s0733g00010,VIT_08s0105g00230,VIT_11s0037g00950,VIT_18s0001g05890,VIT_17s0000g01280,VIT_15s0048g01740,VIT_09s0002g07110 | 3.290270015 | 0.8 | 7.730064482 | 98.60916747 | 1 |
| CARPO All peaks | TRUE | 0.000624338 | 2283 | 6892 | 1011 | 0.146691817 | 0.442838371 | GO:0080090 | GO:BP | regulation of primary metabolic process | 17259 | 20968 | c(GO:0019222, GO:0044238) | VIT_05s0051g00450,VIT_00s0586g00010,VIT_09s0096g00990,VIT_00s0396g00050,VIT_08s0105g00230,VIT_18s0001g05890,VIT_17s0000g01280,VIT_15s0048g01740,VIT_02s0025g04460,VIT_14s0083g00700 | 3.204580497 | 0.8 | 7.717858864 | 156.8328307 | 1 |
| CARPO All peaks | TRUE | 0.000688105 | 347 | 6892 | 180 | 0.026117237 | 0.518731988 | GO:0009755 | GO:BP | hormone-mediated signaling pathway | 17259 | 4123 | c(GO:0007165, GO:0032870) | VIT_05s0062g01120,VIT_05s0020g03310,VIT_14s0030g00440,VIT_17s0000g00320,VIT_08s0007g05740,VIT_13s0047g00250,VIT_08s0007g08300,VIT_09s0002g03410,VIT_05s0077g01390,VIT_11s0016g04490 | 3.162344998 | 0.8 | 6.32491286 | 91.11028114 | 1 |
| CARPO All peaks | TRUE | 0.000945191 | 2512 | 6892 | 1104 | 0.160185723 | 0.439490446 | GO:0060255 | GO:BP | regulation of macromolecule metabolic process | 17259 | 17102 | c(GO:0019222, GO:0043170) | VIT_05s0051g00450,VIT_00s0586g00010,VIT_09s0096g00990,VIT_00s0396g00050,VIT_08s0105g00230,VIT_18s0001g05890,VIT_17s0000g01280,VIT_15s0048g01740,VIT_02s0025g04460,VIT_14s0083g00700 | 3.024480267 | 0.8 | 7.78853601 | 141.7492227 | 1 |
| CARPO All peaks | TRUE | 0.001102629 | 2601 | 6892 | 1140 | 0.16540917 | 0.438292964 | GO:0019222 | GO:BP | regulation of metabolic process | 17259 | 6508 | c(GO:0008152, GO:0050789) | VIT_05s0051g00450,VIT_00s0586g00010,VIT_09s0096g00990,VIT_00s0396g00050,VIT_08s0105g00230,VIT_18s0001g05890,VIT_17s0000g01280,VIT_15s0048g01740,VIT_02s0025g04460,VIT_14s0083g00700 | 2.957570492 | 0.8 | 7.814279043 | 100.415611 | 1 |
| CARPO All peaks | TRUE | 0.001499994 | 2367 | 6892 | 1041 | 0.151044689 | 0.439797212 | GO:0031323 | GO:BP | regulation of cellular metabolic process | 17259 | 8293 | c(GO:0019222, GO:0044237, GO:0050794) | VIT_05s0051g00450,VIT_00s0586g00010,VIT_00s0396g00050,VIT_08s0105g00230,VIT_18s0001g05890,VIT_17s0000g01280,VIT_15s0048g01740,VIT_02s0025g04460,VIT_14s0083g00700,VIT_04s0023g01490 | 2.823910487 | 0.8 | 7.744575096 | 107.3799773 | 1 |
| CARPO All peaks | TRUE | 0.001747505 | 2427 | 6892 | 1065 | 0.154526988 | 0.43881335 | GO:0019438 | GO:BP | aromatic compound biosynthetic process | 17259 | 6708 | c(GO:0006725, GO:0044249) | VIT_05s0051g00450,VIT_00s0586g00010,VIT_04s0079g00620,VIT_00s0733g00010,VIT_08s0105g00230,VIT_11s0037g00950,VIT_18s0001g05890,VIT_17s0000g01280,VIT_15s0048g01740,VIT_09s0002g07110 | 2.757581563 | 0.8 | 7.763083864 | 101.1959321 | 1 |
| CARPO All peaks | TRUE | 0.002572478 | 2531 | 6892 | 1106 | 0.160475914 | 0.43698143 | GO:1901362 | GO:BP | organic cyclic compound biosynthetic process | 17259 | 24654 | c(GO:1901360, GO:1901576) | VIT_05s0051g00450,VIT_00s0586g00010,VIT_04s0079g00620,VIT_00s0733g00010,VIT_08s0105g00230,VIT_11s0037g00950,VIT_18s0001g05890,VIT_17s0000g01280,VIT_15s0048g01740,VIT_09s0002g07110 | 2.589648322 | 0.8 | 7.794107461 | 171.2141497 | 1 |
| CARPO All peaks | TRUE | 0.012886416 | 171 | 6892 | 93 | 0.013493906 | 0.543859649 | GO:0098771 | GO:BP | inorganic ion homeostasis | 17259 | 22204 | GO:0050801 | VIT_19s0014g02740,VIT_08s0007g02240,VIT_02s0025g03360,VIT_19s0015g00320,VIT_18s0001g13410,VIT_05s0020g02160,VIT_05s0020g01150,VIT_02s0488g00010,VIT_05s0020g00440,VIT_00s0577g00040 | 1.889867853 | 0.8 | 5.801677573 | 161.6552155 | 1 |
| CARPO All peaks | TRUE | 0.013223918 | 1489 | 6892 | 663 | 0.096198491 | 0.445265279 | GO:0055085 | GO:BP | transmembrane transport | 17259 | 16858 | c(GO:0006810, GO:0009987) | VIT_00s0438g00010,VIT_00s0733g00010,VIT_02s0012g01160,VIT_18s0122g00090,VIT_07s0031g02960,VIT_05s0062g01120,VIT_08s0007g02240,VIT_18s0001g08270,VIT_02s0025g03360,VIT_17s0119g00080 | 1.87863985 | 0.8 | 7.401855632 | 140.7972309 | 1 |
| CARPO All peaks | TRUE | 0.013366354 | 376 | 6892 | 186 | 0.026987812 | 0.494680851 | GO:0032870 | GO:BP | cellular response to hormone stimulus | 17259 | 9113 | c(GO:0009725, GO:0071310, GO:0071495) | VIT_05s0062g01120,VIT_05s0020g03310,VIT_14s0030g00440,VIT_17s0000g00320,VIT_08s0007g05740,VIT_13s0047g00250,VIT_08s0007g08300,VIT_09s0002g03410,VIT_05s0077g01390,VIT_11s0016g04490 | 1.873987026 | 0.8 | 6.384259261 | 110.579294 | 1 |
| CARPO All peaks | TRUE | 0.013682453 | 1222 | 6892 | 550 | 0.07980267 | 0.450081833 | GO:0032502 | GO:BP | developmental process | 17259 | 8845 | GO:0008150 | VIT_05s0051g00450,VIT_14s0060g02140,VIT_18s0001g09420,VIT_01s0011g04760,VIT_15s0048g01740,VIT_05s0094g01530,VIT_05s0020g03310,VIT_01s0011g00100,VIT_10s0003g00870,VIT_08s0007g03870 | 1.863836027 | 0.8 | 7.255741072 | 109.5336637 | 1 |
| CARPO All peaks | TRUE | 0.013682453 | 178 | 6892 | 96 | 0.013929193 | 0.539325843 | GO:0050801 | GO:BP | ion homeostasis | 17259 | 15396 | GO:0048878 | VIT_19s0014g02740,VIT_08s0007g02240,VIT_02s0025g03360,VIT_19s0015g00320,VIT_18s0001g13410,VIT_05s0020g02160,VIT_05s0020g01150,VIT_02s0488g00010,VIT_05s0020g00440,VIT_00s0577g00040 | 1.863836027 | 0.8 | 5.831341763 | 135.0930832 | 1 |
| CARPO All peaks | TRUE | 0.014735434 | 1119 | 6892 | 506 | 0.073418456 | 0.452189455 | GO:0023052 | GO:BP | signaling | 17259 | 7683 | GO:0008150 | VIT_18s0001g01320,VIT_05s0062g01120,VIT_06s0004g01140,VIT_17s0000g06970,VIT_12s0121g00240,VIT_17s0000g04420,VIT_17s0000g09170,VIT_05s0020g03310,VIT_08s0007g03870,VIT_11s0016g03000 | 1.831637081 | 0.8 | 7.190635537 | 104.9999978 | 1 |
| CARPO All peaks | TRUE | 0.016289848 | 380 | 6892 | 187 | 0.027132908 | 0.492105263 | GO:0071495 | GO:BP | cellular response to endogenous stimulus | 17259 | 19600 | GO:0009719 | VIT_05s0062g01120,VIT_05s0020g03310,VIT_14s0030g00440,VIT_17s0000g00320,VIT_08s0007g05740,VIT_13s0047g00250,VIT_08s0007g08300,VIT_09s0002g03410,VIT_05s0077g01390,VIT_11s0016g04490 | 1.788082968 | 0.8 | 6.392083532 | 151.495434 | 1 |
| CARPO All peaks | TRUE | 0.017256214 | 1114 | 6892 | 503 | 0.072983169 | 0.451526032 | GO:0007165 | GO:BP | signal transduction | 17259 | 3085 | c(GO:0007154, GO:0009987, GO:0023052, GO:0050794, GO:00517 | VIT_18s0001g01320,VIT_05s0062g01120,VIT_06s0004g01140,VIT_17s0000g06970,VIT_12s0121g00240,VIT_17s0000g04420,VIT_17s0000g09170,VIT_05s0020g03310,VIT_08s0007g03870,VIT_11s0016g03000 | 1.763054491 | 0.8 | 7.187324351 | 87.06041433 | 1 |
| CARPO All peaks | TRUE | 0.020013989 | 167 | 6892 | 90 | 0.013058619 | 0.538922156 | GO:0055080 | GO:BP | cation homeostasis | 17259 | 16854 | GO:0050801 | VIT_19s0014g02740,VIT_08s0007g02240,VIT_02s0025g03360,VIT_19s0015g00320,VIT_18s0001g13410,VIT_05s0020g02160,VIT_05s0020g01150,VIT_02s0488g00010,VIT_05s0020g00440,VIT_00s0577g00040 | 1.698666341 | 0.8 | 5.78417648 | 140.7816245 | 1 |
| CARPO All peaks | TRUE | 0.020589891 | 1325 | 6892 | 591 | 0.085751596 | 0.446037736 | GO:0007154 | GO:BP | cell communication | 17259 | 3074 | GO:0009987 | VIT_18s0001g01320,VIT_05s0062g01120,VIT_06s0004g01140,VIT_17s0000g06970,VIT_12s0121g00240,VIT_05s0049g01320,VIT_17s0000g04420,VIT_17s0000g09170,VIT_05s0020g03310,VIT_08s0007g03870 | 1.686345959 | 0.8 | 7.315574903 | 87.01749667 | 1 |
| CARPO All peaks | TRUE | 0.022781952 | 266 | 6892 | 135 | 0.019587928 | 0.507518797 | GO:0033993 | GO:BP | response to lipid | 17259 | 9672 | GO:0010033 | VIT_08s0032g00440,VIT_03s0038g04390,VIT_17s0000g01830,VIT_14s0030g00440,VIT_17s0000g00320,VIT_08s0105g00570,VIT_08s0007g08300,VIT_05s0077g01390,VIT_18s0001g00360,VIT_01s0011g00410 | 1.642409067 | 0.8 | 6.128362827 | 112.7602917 | 1 |
| CARPO All peaks | TRUE | 0.023296206 | 64 | 6892 | 40 | 0.005803831 | 625 | GO:0048437 | GO:BP | floral organ development | 17259 | 14837 | c(GO:0003006, GO:0009908, GO:0099402) | VIT_14s0030g00440,VIT_17s0000g00320,VIT_15s0046g00330,VIT_01s0010g03900,VIT_07s0104g00930,VIT_16s0022g02380,VIT_18s0041g01880,VIT_14s0083g01050,VIT_12s0142g00360,VIT_06s0080g00360 | 1.632714806 | 0.8 | 5.075022777 | 132.9120856 | 1 |
| CARPO All peaks | TRUE | 0.02930433 | 382 | 6892 | 186 | 0.026987812 | 0.486910995 | GO:0048608 | GO:BP | reproductive structure development | 17259 | 14973 | c(GO:0003006, GO:0048856, GO:0061458) | VIT_01s0011g00100,VIT_18s0001g14610,VIT_14s0030g00440,VIT_17s0000g00320,VIT_08s0105g00570,VIT_06s0080g00290,VIT_06s0004g00570,VIT_11s0037g00250,VIT_13s0064g00570,VIT_10s0003g01430 | 1.533068203 | 0.8 | 6.395964836 | 133.442704 | 1 |
| CARPO All peaks | TRUE | 0.035709225 | 67 | 6892 | 41 | 0.005948926 | 0.611940299 | GO:0055067 | GO:BP | monovalent inorganic cation homeostasis | 17259 | 16842 | c(GO:0055080, GO:0098771) | VIT_02s0025g03360,VIT_19s0015g00320,VIT_18s0001g13410,VIT_05s0020g01150,VIT_02s0488g00010,VIT_05s0020g00440,VIT_00s0577g00040,VIT_09s0002g02260,VIT_11s0149g00210,VIT_15s0021g01660 | 1.447219571 | 0.8 | 5.108893738 | 140.7348052 | 1 |
| CARPO All peaks | TRUE | 0.04108762 | 874 | 6892 | 397 | 0.057603018 | 0.45423341 | GO:0010033 | GO:BP | response to organic substance | 17259 | 4328 | GO:0042221 | VIT_08s0032g00440,VIT_00s0733g00010,VIT_05s0062g01120,VIT_02s0154g00010,VIT_17s0000g08450,VIT_05s0020g03310,VIT_03s0038g04390,VIT_17s0000g01830,VIT_14s0030g00440,VIT_17s0000g00320 | 1.386289012 | 0.8 | 7.007925447 | 91.91011033 | 1 |
| CARPO All peaks | TRUE | 0.049177277 | 24 | 6892 | 18 | 0.002611724 | 0.75 | GO:0048466 | GO:BP | androecium development | 17259 | 14866 | GO:0048438 | VIT_15s0046g00330,VIT_06s0080g00360,VIT_07s0031g01180,VIT_13s0067g01760,VIT_08s0007g07750,VIT_18s0001g01760,VIT_14s0068g01570,VIT_08s0007g07760,VIT_04s0023g02650,VIT_12s0059g02740 | 1.308235521 | 0.8 | 4.349810687 | 133.0252322 | 1 |
| CARPO All peaks | TRUE | 0.049177277 | 24 | 6892 | 18 | 0.002611724 | 0.75 | GO:0048443 | GO:BP | stamen development | 17259 | 14843 | c(GO:0048437, GO:0048466, GO:0048827) | VIT_15s0046g00330,VIT_06s0080g00360,VIT_07s0031g01180,VIT_13s0067g01760,VIT_08s0007g07750,VIT_18s0001g01760,VIT_14s0068g01570,VIT_08s0007g07760,VIT_04s0023g02650,VIT_12s0059g02740 | 1.308235521 | 0.8 | 4.349810687 | 132.9354952 | 1 |
| CARPO All peaks | TRUE | 0.049177277 | 475 | 6892 | 225 | 0.032646547 | 0.473684211 | GO:0009791 | GO:BP | post-embryonic development | 17259 | 4153 | c(GO:0007275, GO:0032501) | VIT_01s0011g00100,VIT_18s0001g14610,VIT_14s0030g00440,VIT_17s0000g00320,VIT_08s0105g00570,VIT_06s0080g00290,VIT_06s0004g00570,VIT_11s0037g00250,VIT_13s0064g00570,VIT_10s0003g01430 | 1.308235521 | 0.8 | 6.557072903 | 91.22732932 | 1 |
| CARPO HCT | TRUE | 0.011558146 | 677 | 1240 | 79 | 0.063709677 | 0.116691285 | GO:0009628 | GO:BP | response to abiotic stimulus | 17259 | 4016 | GO:0050896 | VIT_19s0177g00030,VIT_19s0140g00120,VIT_08s0007g01360,VIT_05s0077g01820,VIT_05s0020g01090,VIT_07s0031g01530,VIT_15s0046g02110,VIT_02s0154g00260,VIT_09s0002g02680,VIT_06s0004g06030 | 1.937111818 | 0.8 | 6.819079358 | 90.69280932 | 1 |
| CARPO HCT | TRUE | 0.011558146 | 646 | 1240 | 76 | 0.061290323 | 0.117647059 | GO:0009719 | GO:BP | response to endogenous stimulus | 17259 | 4087 | GO:0050896 | VIT_19s0177g00030,VIT_01s0146g00210,VIT_07s0104g00930,VIT_15s0048g00530,VIT_08s0058g01160,VIT_11s0016g00710,VIT_01s0146g00180,VIT_09s0002g00890,VIT_00s0179g00090,VIT_05s0077g01480 | 1.937111818 | 0.8 | 6.784422996 | 90.96982333 | 1 |
| CARPO HCT | TRUE | 0.011558146 | 642 | 1240 | 76 | 0.061290323 | 0.118380062 | GO:0009725 | GO:BP | response to hormone | 17259 | 4093 | c(GO:0009719, GO:0010033) | VIT_19s0177g00030,VIT_01s0146g00210,VIT_07s0104g00930,VIT_15s0048g00530,VIT_08s0058g01160,VIT_11s0016g00710,VIT_01s0146g00180,VIT_09s0002g00890,VIT_00s0179g00090,VIT_05s0077g01480 | 1.937111818 | 0.8 | 6.779830517 | 90.99323297 | 1 |
| CARPO HCT | TRUE | 0.019406618 | 37 | 1240 | 11 | 0.008870968 | 0.297297297 | GO:0009611 | GO:BP | response to wounding | 17259 | 4003 | GO:0006950 | VIT_11s0016g00710,VIT_07s0031g01530,VIT_09s0002g00890,VIT_00s0179g00090,VIT_10s0003g03810,VIT_01s0011g05560,VIT_18s0001g08310,VIT_18s0001g09420,VIT_02s0025g04720,VIT_02s0025g00360 | 1.712050143 | 0.8 | 4.669864626 | 90.64208845 | 1 |
| CARPO HCT | TRUE | 0.019406618 | 347 | 1240 | 46 | 0.037096774 | 0.132564841 | GO:0009755 | GO:BP | hormone-mediated signaling pathway | 17259 | 4123 | c(GO:0007165, GO:0032870) | VIT_07s0104g00930,VIT_11s0016g00710,VIT_09s0002g00890,VIT_00s0179g00090,VIT_05s0077g01480,VIT_10s0003g03810,VIT_09s0002g03410,VIT_07s0104g01320,VIT_01s0011g05560,VIT_11s0052g00870 | 1.712050143 | 0.8 | 6.32491286 | 91.11028114 | 1 |
| CARPO HCT | TRUE | 0.019406618 | 636 | 1240 | 73 | 0.058870968 | 0.114779874 | GO:1901700 | GO:BP | response to oxygen-containing compound | 17259 | 24953 | GO:0042221 | VIT_14s0068g02330,VIT_05s0077g01820,VIT_03s0063g00370,VIT_07s0104g00930,VIT_11s0016g00710,VIT_15s0046g02110,VIT_09s0002g00890,VIT_02s0154g00260,VIT_00s0179g00090,VIT_10s0003g03810 | 1.712050143 | 0.8 | 6.772887868 | 172.3807298 | 1 |
| CARPO HCT | TRUE | 0.029412943 | 874 | 1240 | 93 | 75 | 0.106407323 | GO:0010033 | GO:BP | response to organic substance | 17259 | 4328 | GO:0042221 | VIT_19s0177g00030,VIT_01s0146g00210,VIT_07s0104g00930,VIT_15s0048g00530,VIT_08s0058g01160,VIT_11s0016g00710,VIT_01s0146g00180,VIT_09s0002g00890,VIT_00s0179g00090,VIT_05s0077g01480 | 1.531461526 | 0.8 | 7.007925447 | 91.91011033 | 1 |
| CARPO HCT | TRUE | 0.032545709 | 17 | 1240 | 7 | 0.005645161 | 0.411764706 | GO:0010114 | GO:BP | response to red light | 17259 | 4408 | GO:0009639 | VIT_05s0020g01090,VIT_02s0025g00880,VIT_01s0011g03160,VIT_15s0046g01510,VIT_18s0001g05690,VIT_07s0031g00450,VIT_14s0068g01350 | 1.487506262 | 0.8 | 4.094840223 | 92.22223879 | 1 |
| CARPO HCT | TRUE | 0.041573269 | 376 | 1240 | 47 | 0.037903226 | 125 | GO:0032870 | GO:BP | cellular response to hormone stimulus | 17259 | 9113 | c(GO:0009725, GO:0071310, GO:0071495) | VIT_07s0104g00930,VIT_11s0016g00710,VIT_09s0002g00890,VIT_00s0179g00090,VIT_05s0077g01480,VIT_10s0003g03810,VIT_09s0002g03410,VIT_07s0104g01320,VIT_01s0011g05560,VIT_11s0052g00870 | 1.38118582 | 0.8 | 6.384259261 | 110.579294 | 1 |
| CARPO HCT | TRUE | 0.048024924 | 380 | 1240 | 47 | 0.037903226 | 0.123684211 | GO:0071495 | GO:BP | cellular response to endogenous stimulus | 17259 | 19600 | GO:0009719 | VIT_07s0104g00930,VIT_11s0016g00710,VIT_09s0002g00890,VIT_00s0179g00090,VIT_05s0077g01480,VIT_10s0003g03810,VIT_09s0002g03410,VIT_07s0104g01320,VIT_01s0011g05560,VIT_11s0052g00870 | 1.318533313 | 0.8 | 6.392083532 | 151.495434 | 1 |
| CARPO All peaks | TRUE | 6.86238651805287E-06 | 2204 | 6900 | 999 | 0.144782609 | 0.453266788 | GO:0005886 | GO:CC | plasma membrane | 17349 | 522 | c(GO:0016020, GO:0071944) | VIT_08s0032g00440,VIT_18s0001g01320,VIT_00s0733g00010,VIT_02s0012g01160,VIT_07s0031g02960,VIT_05s0062g01120,VIT_03s0017g01410,VIT_18s0001g08270,VIT_02s0025g03360,VIT_11s0103g00010 | 5.163524824 | 0.8 | 7.691820271 | 55.75501733 | 2 |
| CARPO All peaks | TRUE | 2.06062833510669E-05 | 2670 | 6900 | 1187 | 0.172028986 | 0.444569288 | GO:0071944 | GO:CC | cell periphery | 17349 | 3147 | GO:0110165 | VIT_08s0032g00440,VIT_18s0001g01320,VIT_00s0733g00010,VIT_02s0012g01160,VIT_07s0031g02960,VIT_05s0062g01120,VIT_12s0134g00160,VIT_03s0017g01410,VIT_18s0001g08270,VIT_02s0025g03360 | 4.686000333 | 0.8 | 7.833638024 | 65.99869292 | 2 |
| CARPO All peaks | TRUE | 0.042615386 | 426 | 6900 | 204 | 0.029565217 | 0.478873239 | GO:0009506 | GO:CC | plasmodesma | 17349 | 723 | c(GO:0005911, GO:0055044) | VIT_08s0032g00440,VIT_11s0037g00770,VIT_14s0068g00110,VIT_18s0001g14610,VIT_00s0225g00100,VIT_16s0022g01430,VIT_06s0004g03260,VIT_00s0264g00030,VIT_08s0007g01700,VIT_18s0072g00160 | 1.370433578 | 0.8 | 6.476571839 | 56.53939021 | 2 |
| CARPO All peaks | TRUE | 0.042615386 | 426 | 6900 | 204 | 0.029565217 | 0.478873239 | GO:0005911 | GO:CC | cell-cell junction | 17349 | 544 | GO:0070161 | VIT_08s0032g00440,VIT_11s0037g00770,VIT_14s0068g00110,VIT_18s0001g14610,VIT_00s0225g00100,VIT_16s0022g01430,VIT_06s0004g03260,VIT_00s0264g00030,VIT_08s0007g01700,VIT_18s0072g00160 | 1.370433578 | 0.8 | 6.476571839 | 55.84086909 | 2 |
| CARPO All peaks | TRUE | 0.042615386 | 426 | 6900 | 204 | 0.029565217 | 0.478873239 | GO:0070161 | GO:CC | anchoring junction | 17349 | 2780 | GO:0030054 | VIT_08s0032g00440,VIT_11s0037g00770,VIT_14s0068g00110,VIT_18s0001g14610,VIT_00s0225g00100,VIT_16s0022g01430,VIT_06s0004g03260,VIT_00s0264g00030,VIT_08s0007g01700,VIT_18s0072g00160 | 1.370433578 | 0.8 | 6.476571839 | 64.56652951 | 2 |
| CARPO All peaks | TRUE | 0.042615386 | 428 | 6900 | 205 | 0.029710145 | 0.478971963 | GO:0030054 | GO:CC | cell junction | 17349 | 1035 | GO:0110165 | VIT_08s0032g00440,VIT_11s0037g00770,VIT_14s0068g00110,VIT_18s0001g14610,VIT_00s0225g00100,VIT_16s0022g01430,VIT_06s0004g03260,VIT_00s0264g00030,VIT_08s0007g01700,VIT_18s0072g00160 | 1.370433578 | 0.8 | 6.480035015 | 57.75692422 | 2 |
| CARPO All peaks | TRUE | 0.042615386 | 426 | 6900 | 204 | 0.029565217 | 0.478873239 | GO:0055044 | GO:CC | symplast | 17349 | 2582 | GO:0110165 | VIT_08s0032g00440,VIT_11s0037g00770,VIT_14s0068g00110,VIT_18s0001g14610,VIT_00s0225g00100,VIT_16s0022g01430,VIT_06s0004g03260,VIT_00s0264g00030,VIT_08s0007g01700,VIT_18s0072g00160 | 1.370433578 | 0.8 | 6.476571839 | 63.79386369 | 2 |
| CARPO All peaks | TRUE | 2.89667643233272E-13 | 812 | 7595 | 437 | 0.057537854 | 0.53817734 | GO:0003700 | GO:MF | DNA-binding transcription factor activity | 19061 | 428 | GO:0140110 | VIT_05s0051g00450,VIT_08s0105g00230,VIT_18s0001g05890,VIT_17s0000g01280,VIT_02s0025g04460,VIT_14s0083g00700,VIT_01s0011g00100,VIT_10s0003g00870,VIT_18s0072g00260,VIT_03s0038g00860 | 12.53810001 | 0.8 | 6.953521349 | 3.666066962 | 3 |
| CARPO All peaks | TRUE | 4.5353599265561E-13 | 964 | 7595 | 505 | 0.066491113 | 0.523858921 | GO:0140110 | GO:MF | transcription regulator activity | 19061 | 11795 | GO:0003674 | VIT_05s0051g00450,VIT_08s0105g00230,VIT_18s0001g05890,VIT_17s0000g01280,VIT_02s0025g04460,VIT_14s0083g00700,VIT_01s0011g00100,VIT_10s0003g00870,VIT_18s0072g00260,VIT_03s0038g00860 | 12.34338824 | 0.8 | 7.080393416 | 48.01778397 | 3 |
| CARPO All peaks | TRUE | 1.23524561474112E-08 | 818 | 7595 | 418 | 0.055036208 | 0.511002445 | GO:0043565 | GO:MF | sequence-specific DNA binding | 19061 | 6871 | GO:0003677 | VIT_05s0051g00450,VIT_08s0105g00230,VIT_17s0000g01280,VIT_01s0011g04760,VIT_09s0002g07110,VIT_06s0061g00760,VIT_11s0149g00140,VIT_01s0011g00100,VIT_10s0003g00870,VIT_17s0000g00320 | 7.908246679 | 0.8 | 6.958964711 | 28.80533965 | 3 |
| CARPO All peaks | TRUE | 3.8849054371407E-05 | 2015 | 7595 | 913 | 0.120210665 | 0.453101737 | GO:0003677 | GO:MF | DNA binding | 19061 | 408 | GO:0003676 | VIT_05s0051g00450,VIT_04s0079g00620,VIT_08s0105g00230,VIT_18s0001g05890,VIT_17s0000g01280,VIT_01s0011g04760,VIT_09s0002g07110,VIT_02s0025g04460,VIT_18s0001g01350,VIT_06s0061g00760 | 4.410619548 | 0.8 | 7.625530782 | 3.588031039 | 3 |
| CARPO All peaks | TRUE | 0.03143662 | 334 | 7595 | 167 | 0.02198815 | 0.5 | GO:0001067 | GO:MF | regulatory region nucleic acid binding | 19061 | 182 | GO:0003676 | VIT_08s0105g00230,VIT_01s0011g00100,VIT_11s0016g04130,VIT_01s0026g02430,VIT_16s0098g01830,VIT_15s0046g00320,VIT_02s0012g01320,VIT_01s0026g02460,VIT_04s0008g01820,VIT_04s0008g06130 | 1.502564152 | 0.8 | 6.296680276 | 2.706225106 | 3 |
| CARPO All peaks | TRUE | 0.03143662 | 334 | 7595 | 167 | 0.02198815 | 0.5 | GO:0000976 | GO:MF | transcription regulatory region sequence-specific DNA binding | 19061 | 128 | c(GO:0001067, GO:1990837) | VIT_08s0105g00230,VIT_01s0011g00100,VIT_11s0016g04130,VIT_01s0026g02430,VIT_16s0098g01830,VIT_15s0046g00320,VIT_02s0012g01320,VIT_01s0026g02460,VIT_04s0008g01820,VIT_04s0008g06130 | 1.502564152 | 0.8 | 6.296680276 | 2.495528113 | 3 |
| CARPO All peaks | TRUE | 0.048834343 | 360 | 7595 | 177 | 0.023304806 | 0.491666667 | GO:1990837 | GO:MF | sequence-specific double-stranded DNA binding | 19061 | 12209 | c(GO:0003690, GO:0043565) | VIT_08s0105g00230,VIT_11s0149g00140,VIT_01s0011g00100,VIT_11s0016g04130,VIT_01s0026g02430,VIT_16s0098g01830,VIT_15s0046g00320,VIT_02s0012g01320,VIT_01s0026g02460,VIT_04s0008g01820 | 1.311274648 | 0.8 | 6.352106948 | 49.63312758 | 3 |
| CARPO All peaks | TRUE | 0.048834343 | 1414 | 7595 | 627 | 0.082554312 | 0.443422914 | GO:0022857 | GO:MF | transmembrane transporter activity | 19061 | 4706 | GO:0005215 | VIT_00s0438g00010,VIT_00s0733g00010,VIT_02s0012g01160,VIT_18s0122g00090,VIT_07s0031g02960,VIT_05s0062g01120,VIT_08s0007g02240,VIT_18s0001g08270,VIT_02s0025g03360,VIT_17s0119g00080 | 1.311274648 | 0.8 | 7.363642512 | 20.35795095 | 3 |
| VHCT | TRUE | 0.008145049 | 49 | 26 | 4 | 0.153846154 | 0.081632653 | KEGG:01210 | KEGG | 2-Oxocarboxylic acid metabolism | 4166 | 184 | KEGG:00000 | VIT_18s0001g08430,VIT_17s0000g07790,VIT_04s0008g03770,VIT_09s0002g08670 | 2.089106312 | 0.8 | 4.87756011 | 198.6045907 | 1 |
| VHCT | TRUE | 0.023923655 | 44 | 26 | 3 | 0.115384615 | 0.068181818 | KEGG:00250 | KEGG | Alanine, aspartate and glutamate metabolism | 4166 | 26 | KEGG:00000 | VIT_18s0001g09910,VIT_04s0008g03770,VIT_16s0098g00290 | 1.621172461 | 0.8 | 4.797979429 | 197.9870177 | 1 |
| VHCT | TRUE | 0.023923655 | 10 | 26 | 2 | 0.076923077 | 0.2 | KEGG:00966 | KEGG | Glucosinolate biosynthesis | 4166 | 151 | KEGG:00000 | VIT_18s0001g08430,VIT_17s0000g07790 | 1.621172461 | 0.8 | 3.702500759 | 198.4756039 | 1 |
| VHCT | TRUE | 0.023923655 | 231 | 26 | 6 | 0.230769231 | 0.025974026 | KEGG:04075 | KEGG | Plant hormone signal transduction | 4166 | 242 | KEGG:00000 | VIT_15s0048g00530,VIT_11s0016g00710,VIT_18s0001g06310,VIT_01s0011g05560,VIT_11s0016g04490,VIT_13s0019g02200 | 1.621172461 | 0.8 | 6.02405119 | 198.8312946 | 1 |

**Tab. S3.** Lists of primers used.

| Gene name | Gene ID | Purpose | Sequence 5'-3' |
|---|---|---|---|
| ACTIN | Solyc03g078400 | qPCR | FOR: GCCGCATGCCATTCTTCGTT<br>REV: TCCCGTTCAGCAGTGGTGG |
| SlACS4 | Solyc05g050010 | qPCR | FOR: AGATCGCACTTGCAAGGATTC<br>REV: TCCCGTTCAGCAGTGGTGG |
| SlPG2a | Solyc10g080210 | qPCR | FOR: TCAAGGGCACAAGTGCAACAA<br>REV: TGCACGTAGCCTCTGATGGT |
| SlPSY1 | Solyc03g031860 | qPCR | FOR: ACAGGCAGGTCTATCCGATG<br>REV: AGCTCAATTCTGTCACGCCTT |
| SlSGR1 | Solyc08g080090 | qPCR | FOR: ACCATCCAACAAGGCTTGCA<br>REV: GCTGCTTCCACAAACCCTAT |
| VviCARPO | VIT_08s0007g07670 | Gene Isolation (Overexpressing Plants) | FOR: CACCATGGACAACCCGCAATCCAC<br>REV: TCATCCTTGAAATGGGAAATAAG |
| | | Gene Isolation (Repressing Plants) | REV: TTAAGCGAAACCCAAACGGAGTTCTAGATCCAGATCGAGTCCTTGAAATGGGAAATAAG |
| | | qPCR | FOR: CTTCCACACTCGGTCATCTC<br>REV: GGATGTCGTTGGATTGGCTG |
| | | Promoter Isolation | FOR ext: TGGCCATTCATGCGAGGTAT<br>FOR int: CCCAAGCTTGGGTGTTGCCAATCGAATTGATGG |
| | | | REV ext: TCGACATCGGCGATGATGGA<br>REV int: GGACTAGTCCGGCTGTCGCTGAAAAATTATG |
| VviMyb14 | VIT_07s0005g03340 | qPCR | FOR: TCTGAGGCCGGATATCAAAC<br>REV: GGGACGCATCAAGAGAGTG |
| | | Promoter Isolation | FOR: CACCGGCTTCACCAATCATAGAGCTTA<br>REV: TTTTTCTTTTCTACGTAAGGA |
| VviMYBA1 | VIT_02s0033g00380 | qPCR | FOR: TAGTCACCACTTCAAAAAGG<br>REV: GAATGTGTTTGGGGTTTATC |
| | | Promoter Isolation | FOR ext: CCCTTCATTAACATCAAAAGGTC<br>FOR int: CACCTGGAGAAAATGATGCAACGA<br>REV ext: GAACCTCCTTTTTGAAGTGGTGACT<br>REV int: CGAGTCAACTCAACACAAGAGA |
| VviNAC03 | VIT_00s0375g00040 | qPCR | FOR: TGCCCTGCTTCTCCGATATG<br>REV: CTGGCATTCCTCCAAATATGG |
| VviNAC33 | VIT_19s0027g00230 | qPCR | D'Incà et al. 2021 |
| VviSGR1 | VIT_02s0025g04660 | qPCR | D'Incà et al. 2021 |
| | | Promoter Isolation | D'Incà et al. 2021 |
| VviUBIQUITIN | VIT_16s0098g01190 | qPCR | FOR: TCTGAGGCTTCGTGGTGGTA<br>REV: AGGCGTGCATAACATTTGCG |
| VviWYRKY16 | VIT_06S0004G07500 | qPCR | FOR: ATAAGTGCACGAACCCAGGA<br>REV: CACATCATGGTTGTGCTTCC |

**Tab. S4.** Number of $T_1$ tomato plants obtained from $T_0$ generation.

| Construct | Line | N° $T_1$ plants |
|---|---|---|
| 35S::*VviCARPO* | #1 | 10 |
| | #3 | 14 |
| 35S::*VviNAC03* | #1 | 12 |
| | #12 | 9 |
| 35S::*VviNAC33* | #2 | 15 |
| | #3 | 16 |

**Supplemental Datasets**

**Dataset S1.** List of the highest CARPO co-expression genes.

**Dataset S2. D**AP-Seq_All peaks CARPO.

**Dataset S3.** Differentially expressed genes in stable overexpressing CARPO transgenic plants.

**Dataset S4.** Differentially expressed genes in transient overexpressing CARPO transgenic plants.

**Dataset S5.** High confident targets of CARPO identified by combining DAP-seq data with transcriptomic analysis.

**Dataset S6.** DAP-Seq_All peaks VviNAC03.

**Dataset S7.** DAP-Seq_All peaks VviNAC33.

**Dataset S8.** VviNAC03, VviNAC33 and CARPO filtered DAP-Seq bound genes.

**Dataset S9.** Protein sequences of grapevine, tomato and Arabidopsis and all characterized NACs in any other species.
